# Bacterial IF2’s N-terminal IDR drives cold-induced phase separation and promotes fitness during cold stress

**DOI:** 10.1101/2025.02.02.631968

**Authors:** Aishwarya Ghosh, Kaveendya S. Mallikaarachchi, Kathryn G. Dzurik, Vidhyadhar Nandana, Nathaniel R. Nunez, W. Seth Childers, Jared M. Schrader

## Abstract

Translation initiation factor 2 (IF2) plays an essential role in bacterial cells by delivering the fMet-tRNA^fMet^ to the ribosome pre-initiation complex. IF2 is known to have an N-terminal disordered region which is present across bacterial species, yet its function is not fully understood. Deletion of the IDR in *E. coli* showed no phenotypes at normal growth temperature (37°C); however, this IDR was found to be required for growth at cold temperatures (15°C). Since large IDRs can drive phase separation of various RNA binding proteins into biomolecular condensates, we investigated whether *E. coli* IF2 could phase separate. We discovered that IF2’s N-terminal IDR drives phase separation in *E. coli* and *C. crescentus*, suggesting that IF2 condensation is a conserved property. Finally, using *E. coli*, we found that the IDR strongly drives phase separation in the cold, suggesting IF2 condensates promote fitness during cold stress.

**Highlights:**

- IF2’s IDR promotes phase separation with RNA.
- Cold temperature promotes IF2 condensation with RNA.
- IF2’s IDR promotes fitness during cold shock.

## Introduction

Translation initiation is an essential process for expression of the genetic code whereby the ribosome and initiator tRNA are loaded on the mRNA start codon^1,2^. Translation initiation factor 2 (IF2) plays a key role in translation initiation by delivering fMet-tRNA^fMet^ to the ribosome forming the pre-initiation complex, and releasing fMet-tRNA^fMet^ into the P-site after hydrolyzing its bound GTP, allowing the ribosome to enter translation elongation^3,4^. Despite the important role for IF2 in translation initiation, IF2 contains a small N-terminal subdomain and a large intrinsically disordered region (IDR)^5–7^ which is poorly understood. In addition, *E. coli* cells are known to produce three protein isoforms of IF2^8,9^, with distinct regions of the IDR truncated, yet all isoforms contain the GTP, tRNA, and fMet binding domains and are all known to be capable of translation initiation activity. Initial work on IF2’s IDR established that deletion of the N-terminal IDR led to a cold-sensitive phenotype where *E. coli* cells were unable to grow at 15 °C^10^. Additionally, a drug screen targeting ribosome assembly inhibitors found a novel antibiotic candidate, lamotrigine, which bound to the N-terminal IDR and led to a cold-specific inhibition of *E. coli* growth^11^. Interestingly, the N-terminal IDR also promotes dsDNA break repair^12,13^, suggesting that IF2’s IDR may be involved in cellular functions outside of translation initiation. Despite these studies, the exact molecular and cellular function(s) of the N-terminal IDR have remained poorly understood.

Biomolecular condensates are phase separated membraneless assemblies that typically assemble through the physical process of phase separation^14^. Many RNA processes are spatially localized in biomolecular condensates^14^, including rRNA transcription and assembly which occur in the nucleolus^15^, mRNA storage in p-granules^16^, and RNA decay and storage which occur in P-bodies^17,18^. While some condensates like nucleoli and p-bodies are constitutively present, other condensates such as Ded1P or poly-A binding protein condense during heat shock^19,20^. Stress granules contain stalled translation initiation complexes^21,22^ and condense under a variety of stresses^23,24^ including heat shock, oxidative stress, viral infection, osmotic stress, UV irradiation, or cold shock. Interestingly, eIF2α is known to be a key regulator of the stress granule formation, in which its phosphorylation leads to a shutdown in translation that triggers stress granule condensation and leads to enhanced fitness under stresses like cold shock^25^. Interestingly, biomolecular condensates have been recently discovered in bacteria^26,27^, and are involved in many in RNA processes^28^, with RNAP condensates corresponding to rRNA transcription^29^, BR-bodies promoting mRNA decay^30^, and Hfq condensates involved in mRNA/sRNA regulation and chromatin compaction^31,32^. Importantly, biomolecular condensate phase separation is often driven by intrinsically disordered regions making multi-valent protein-protein, protein-RNA, and RNA-RNA interactions^14,33,34^. However, no bacterial biomolecular condensates have been discovered whose phase separation is driven by the translation machinery.

To investigate bacterial IF2’s potential for phase separation, we investigated the N-terminal IDR and found that it is present across most bacteria and is predicted to phase separate in a majority of species by DeePhase or fuzdrop algorithms. Purified *E. coli* and *C. crescentus* IF2 phase separate with total RNA at 25°C, and the N-terminal IDR drives phase separation of these proteins, suggesting the ability to drive phase separation is broadly conserved. We found that ribosomes and initiator tRNA were able to prevent IF2 phase separation and dissolve IF2 condensates. We find that cold temperature (4°C) could strongly promote IF2 phase separation which requires the IDR, and that reversible IF2 foci were observed *in vivo* upon cold shock at 4°C. Finally, we found that *E. coli* cells lacking the IDR were sensitive to cold shock, suggesting that IF2 condensates protect *E. coli* against adaptation to cold temperatures. Taken together, IF2 condensates are a conserved environmentally responsive condensate that promotes fitness during adaptation to cold temperature with similar properties to eukaryotic stress granules.

## Materials and Methods

### Raw data deposition

Raw data reported within this manuscript has been uploaded to figshare with DOI: 10.6084/m9.figshare.27624369.

### Plasmid construction

#### *E. coli* plasmid construction

##### pET-MBP-TEV-His-infB-Ec

The *E. coli* IF2 gene was inserted into the pET-MBP-TEV-His vector using SspI and HindIII restriction sites. The forward primer VN102F (GCTCATATGATGACAGATGTAACGATTAAAACG) and the reverse primer VN103R (TTAAAGCTTTTAAGCAATGGTACGTTGGATCTC) were used to amplify the IF2 gene for cloning.

##### pET-GST-TEV-His- ΔNTDinfB-Ec

The *E. coli* ΔNTD_IF2 gene was cloned in the pET-GST-TEV-His vector using SspI and HindIII restriction sites.The forward primer VN104F (GCTAATATTCTGCAGCAAGGCTTCCAGAAG) and the reverse primer VN103R (TTAAAGCTTTTAAGCAATGGTACGTTGGATCTC) were used to amplify the gene for cloning.

##### pET-GST-TEV-His- βinfB-Ec

The *E. coli* beta isoform of IF2 gene is cloned in the pET-GST-TEV-His vector using SspI and HindIII restriction sites. The forward primer VN105F (GCTAATATTAGCAATCAACAAGACGATATGACT) and VN103R (TTAAAGCTTTTAAGCAATGGTACGTTGGATCTC) were used to amplify the gene.

##### pET28a-His-infB-Ec

The *E.coli* IF2 gene is cloned in the pET28a vector using NdeI and EcoRI sites. The forward primer NN1 (NdeI) AAACATATGAGCGACGAGAACGAAAAC and reverse primer NN2 (EcoR11) (AAAGAATTCCTAGTCGAGCTGGCGCTTGATC) is used to amplify the gene.

#### *C. crescentus* IF2 plasmid construction

##### pET28a-His-infB-Cc

The *C. crescentus* IF2 gene is cloned in the pET28a vector using NdeI and EcoRI sites. The forward primer J5_00301_F (NdeI) (CGGCAGCCATATGAGCGACGAGAACGAAAACGGC) and reverse primer J5_00303_R (EcoRI) (CGGAGCTCGAATTCCTAGTCGAGCTGGCGCTTGATCTCTTCG) is used to amplify the gene.

##### pET28a-His-infBΔNTD-Cc

The *C. crescentus* ΔNTD_IF2 gene was cloned by inverse PCR using the pET28a-C.cr IF2 plasmid as template and the following PCR primers AG06_R (ATGGCTGCCGCGCGGCACCA)and AG07_F- (GCCGCGCCGAACAAGGCCGT). The *E. coli* and *C.crescentus’s* IF-2 gene was cloned into pET28a and pET His6 MBP TEV plasmids, and the expressed proteins were purified using His-tag affinity purification

### Bacterial strains used

#### Strains from others

SPA Tagged *E. coli* CloneId:INFB^35^ ORF strain are generated as described (37).

*E. coli* BL21infB::KanR pBAD-infB strains are generated as described (10).

*E. coli* BL21infB::KanR pBAD-infB- ΔNTD strains are generated as described (10)

#### Strain construction

##### JS800

The *E. coli* BL21 strain was transformed with the pET28a-His plasmid, which contains the *E.coli* infB gene cloned using the method described above. The transformation was performed in BL21 cells carrying Kanamycin resistance (KanR), resulting in the generation of E.coli BL21::KanR pET28a-His-infB.

##### JS791

The *E. coli* BL21 strain was transformed with the pET-MBP-TEV-His plasmid, which contains the *E.coli* infB gene cloned using the method described above. The transformation was performed in BL21 cells carrying Kanamycin resistance (KanR), resulting in the generation of E.coli BL21::KanR pET-MBP-TEV-His -infB strains.

##### JS792

The *E. coli* BL21 strain was transformed with the pET-GST-TEV-His plasmid, which contains the *E.coli* infB ΔNTD gene cloned using the method described above. The transformation was performed in BL21 cells carrying Kanamycin resistance (KanR), resulting in the generation of BL21::KanR pET-GST-TEV-His -infB ΔNTD strains.

##### JS793

The *E. coli* BL21 strain was transformed with the pET-GST-TEV-His plasmid, which contains the *E.coli* infB β gene cloned using the method described above. The transformation was performed in BL21 cells carrying Kanamycin resistance (KanR), resulting in the generation of E.coli BL21::KanR pET-GST-TEV-His -infB β form.

##### JS794

The *E. coli* BL21 strain was transformed with the pET28a-His plasmid, which contains the *C.crescentus* infB gene cloned using the method described above. The transformation was performed in BL21 cells carrying Kanamycin resistance (KanR), resulting in the generation of E.coli BL21::KanR pET28a-His-infB.

##### JS795

The *E. coli* BL21 strain was transformed with the pET28a-His plasmid, which contains the *C.crescentus* infB ΔNTD gene cloned using the method described above. The transformation was performed in BL21 cells carrying Kanamycin resistance (KanR), resulting in the generation of *E. coli* BL21::KanR pET28a-His-infB- ΔNTD.

### Protein purification

The pET overexpression plasmid constructs were expressed in BL21 DE3 cells by inducing with 0.5 mM IPTG at 37°C for 3 hours in 2X Luria-Bertani medium. The cells were harvested by centrifugation at 5,000 rpm for 10 minutes and lysed in a lysis buffer containing 20 mM Tris pH 7.4, 500 mM NaCl, 1 mM DTT, 10 mM imidazole, 10% glycerol, 1 mM PMSF, protease inhibitor cocktail [thermo-scientific], and 10 μg/ml DNase I at 4°C using sonication with 10- second pulse-on and 30-second pulse-off cycles at 50% amplitude for 2 minutes and 30 seconds. The lysed cells were centrifuged at 15,000 rpm for 45 minutes, and the supernatant was loaded onto an equilibrated Ni-NTA resin [thermo-scientific]. The protein-bound resin was washed with 10 column volumes each of lysis buffer, chaperone buffer (20 mM Tris pH 7.4, 1 M NaCl, 10% glycerol, 10 mM imidazole, 5 mM KCl, 10 mM MgCl_2_, and 1 mM DTT), and low salt buffer (20 mM Tris pH 7.4, 1 M NaCl, 10% glycerol, 10 mM imidazole, and 1 mM DTT). The proteins were eluted using an elution buffer (20 mM Tris pH 7.4, 150 mM NaCl, 10% glycerol, 250 mM imidazole, and 1 mM DTT), dialyzed into a reaction buffer (20 mM Tris pH 7.4, 75 mM KCl, 10 mM MgCl_2_, and 1 mM DTT), and the concentrated proteins were stored at −80°C.

### Total RNA extraction

For RNA extraction, *E. coli* (DH10β) cells were grown to mid-log phase (OD600 0.4–0.5), and cells were harvested by centrifugation at 7,200xg for 45 seconds. The cell pellet was resuspended in 20 mL of pre-heated (65°C) Trizol and incubated at 65°C for 10 minutes in a water bath. Subsequently, 20 mL of chloroform was added, and the mixture was incubated for 5 minutes at room temperature. The samples were then centrifuged at maximum speed (22,000xg) for 10 minutes at 4°C. The aqueous layer was transferred to a new tube, and 20 mL of chloroform was added, followed by vortexing and centrifugation at 22,000xg for 10 minutes. The upper aqueous layer was collected, and 3 times the volume of ice-cold 100 % ethanol, 1/10th volume of sodium acetate, and 5 μL of gluco-blue were added. The mixture was incubated overnight at −80°C. After 12 hours, the samples were centrifuged at maximum speed for 60 minutes at 4°C. The supernatant was discarded, and the pellet was washed with ice-cold 80% ethanol. After removing the supernatant and drying the pellet, it was resuspended in nuclease-free water. The quality of the extracted RNA was assessed using the Agilent Tape Station platform.

### Turbidity assay

#### 2D phase diagram

A 100 μL reaction volume was set up in a clear bottom 96-well plate using a buffer consisting of 20 mM Tris pH 7.4, 75 mM KCl, 10 mM MgCl2, and 1 mM DTT. The proteins were incubated at various concentrations, ranging from below to above their physiological levels (1.25 μM, 2.5 μM, 5 μM, 10 μM, 20 μM, and 40 μM), either in the absence or presence of a range of *E. coli* total RNA concentrations. Using a plate reader, the turbidity of the solutions was measured by monitoring the optical density at 650 nm (OD650). Reactions were also imaged on a phase contrast microscope (Evos M5000 AMF 5000) to confirm the presence of IF2 droplets.

#### Reversibility of turbidity

500 mM KCl or 60 nM RNase A was added to the samples containing 5 μM of IF2 protein with 5ng/µl of *E. coli* total RNA that were pre-formed for 30 minutes. After incubation, the turbidity was again measured by monitoring OD650.

### *In vitro* phase separation experiments and droplet assays by phase contrast microscopy

#### Protein-RNA phase separation of purified IF2

Proteins were incubated at a final concentration of 5 μM with *E. coli* total RNA (5 ng/μL) in a buffer containing 20 mM Tris (pH 7.4), 75 mM KCl, 10 mM MgCl₂, and 1 mM DTT. The reaction mixture was incubated for 30 minutes at room temperature (25°C). After incubation, the sample was gently pipetted onto a glass slide and covered with a coverslip. Imaging was conducted using an phase contrast microscope. A similar protocol was followed for *C. crescentus*, where proteins were incubated at 5 μM concentration with *C. crescentus* total RNA (50 ng/μL) in the same buffer composition. The mixture was incubated for 30 minutes at room temperature and subsequently imaged as described above.

The *E.coli* MBP-IF2 fusion protein, at its physiological concentration of 20μM was incubated in a buffer containing 20 mM Tris (pH 7.4), 75 mM KCl, 10 mM MgCl_2_, and 1 mM DTT. To trigger the cleavage of the solubilizing MBP tag, a ratio of IF2 to TEV protease was 1:15 added. This mixture was allowed to incubate for 2.5 hours at room temperature. Subsequently, a small portion of the incubated mixture was pipetted onto a glass slide, covered with a coverslip, and imaged under a phase contrast microscope (Evos M5000 AMF5000).

For each experiment, at least five independent images per condition were taken from three experimental replicates to quantify the area of protein-RNA droplets. The images were analyzed using the “analyze particles” function in ImageJ to quantify droplet area across all images. The average of all three replicates was plotted on a bar graph and error bar is represented by standard error. The t-test (one-tailed distribution and two sample unequal variances) is used to estimate the statistical significance.

#### Dissolution of purified IF2-RNA droplets by high-salt or RNase treatments

To assess droplet formation reversibility, the initial droplets were formed using the procedure described above. After confirming droplet formation under the microscope, 500 mM KCl or 60 nM RNase A was added to the reaction and incubated for 40 minutes at room temperature. Following incubation, the mixture was pipetted onto a glass slide, covered with a coverslip, and imaged under a microscope.

#### Addition of tRNA^fMet^ and 70S ribosomes to IF2 droplet reactions

To check if tRNA^fMet^ and 70S ribosome inhibit the droplet formation, proteins were incubated at 5 μM with *E. coli* total RNA (5 ng/μL), along with either 10 μM 70S tightly coupled ribosomes or 5 μM tRNA^fMet^ (gift from O.C. Uhlenbeck), in the buffer composed of 20 mM Tris (pH 7.4), 75 mM KCl, 10 mM MgCl₂, and 1 mM DTT. The reaction was incubated for 30 minutes at room temperature. After incubation, the mixture was transferred onto a glass slide, covered with a coverslip, and imaged using phase contrast microscope.

#### Dissolution of IF2-RNA droplets by tRNA^fMet^ and 70S ribosomes

Droplets were initially formed with IF2 at 5 μM and *E. coli* total RNA (5 ng/μL) in the same buffer conditions as described above. The reaction was incubated for 30 minutes at room temperature, and droplet formation was confirmed under the microscope. Subsequently, 10 μM 70S ribosomes or 5 μM tRNA^fMet^ were added to the same reaction, followed by an additional 30- minute incubation. The mixture was then pipetted onto a glass slide, covered with a coverslip, and imaged under a microscope.

#### Addition of GDP or GTP to IF2-RNA reactions

Proteins were incubated at a final concentration of 5 μM with *E. coli* total RNA (5 ng/μL) and 1mM GTP or GDP added in a buffer containing 20 mM Tris (pH 7.4), 75 mM KCl, 10 mM MgCl₂, and 1 mM DTT. The reaction mixture was incubated for 30 minutes at room temperature (25°C). After incubation, the sample was gently pipetted onto a glass slide and covered with a coverslip. Imaging was conducted using phase contrast microscope.

#### Altering IF2-RNA incubation temperature

IF2-RNA droplets were formed using 5 μM with *E. coli* total RNA (5 ng/μL) in the standard buffer (20 mM Tris, pH 7.4, 75 mM KCl, 10 mM MgCl₂, and 1 mM DTT) and incubated for 30 minutes at different temperatures (4°C, 25°C, and 37°C). After incubation, samples were pipetted onto a glass slide, covered with a coverslip, and imaged using a microscope.

Droplets were formed at 4°C using the protocol described above. Droplet formation was confirmed under the microscope, and the reaction was subsequently incubated at room temperature for 30-40 minutes. After this incubation, the samples were imaged to assess the effect of temperature on droplet reversibility.

### Mant-GTP binding assay

100 µL of reaction mixture contained 1 µM or 0 µM of protein with or without 125 µM of MANT GTP (Thermofisher) in reaction buffer (20 mM Tris (pH 7.4), 75 mM KCl, 10 mM MgCl₂, and 1 mM DTT) in black 96 well plate with clear bottom (corning). The resulting solution at room temperature was excited at 355 nm and the emitted fluorescence values were collected from at 450 nm using Spectramax M2 plate reader. The fluorescence reading at 450 nm from softmax pro software was used to plot the data as a ratio of fluorescence (Protein+MANT GTP/MANT GTP). Cold chase reaction mixtures contained an additional 3 mM GTP.

### IF2 Immunofluorescence

*E. coli* cells harboring the IF-2 gene tagged with a flag epitope and MG1655(control) cells were cultured in M9 minimal media substituted with glucose, MgSo4, and Cacl2 at 37°C with shaking until reaching the log phase (OD600 = 0.5–0.6). Following the growth phase, the bacterial culture was subjected to different fixation conditions:

1. **Cold Incubation and Fixation**: The bacterial culture was incubated at 4°C for 1 hour to simulate cold shock conditions. After incubation, a 2 mL aliquot of the culture was collected and fixed with 4% formaldehyde at 4°C for 4 hours.
2. **Control Fixation**: For the control condition, a 2 mL aliquot of the culture was directly fixed with 4% formaldehyde at 37°C for 30 minutes in a shaker incubator, without prior cold incubation.
3. **Fixation after cold-shock recovery**: For the fixation under conditions where the cells were allowed to recover after cold-shock, the bacterial culture (OD600 = 0.5–0.6) was initially incubated at 4°C for 1 hour, followed by re-incubation at 37°C with shaking for 1 hour. After this temperature shift, the cells were collected and fixed using the same procedure as described above.

#### Cell Preparation

After fixation, cells were pelleted by centrifugation at 6000 rpm in a microcentrifuge for 2 minutes and washed three times with 1X PBST (PBS with 0.1% Tween-20). The cell pellet was gently resuspended in 0.2 mL of GTE buffer (50 mM glucose, 10 mM EDTA, 20 mM Tris-HCl, pH 7.4) containing 750 μg/mL lysozyme and incubated at room temperature for 30 minutes.

#### Slide Preparation

A drop of the treated cell suspension was placed on a glass slide pre-coated with 0.1% w/v poly-L-lysine and allowed to air dry for 20–30 minutes. The slides were then washed three times with 1X PBST and submerged in ice-cold ethanol overnight, stored at 4°C.

#### Immunostaining Procedure

The following day, the slide samples were rehydrated in 1X PBST and blocked with 5% non-fat milk dissolved in 1X PBST for 30 minutes at room temperature with gentle shaking. The slides were then incubated with primary anti-Flag antibody (1:1000 dilution) for 1 hour at room temperature. After incubation, the slides were washed three times with 1X PBST and re-blocked with 5% milk for 30 minutes.

#### Secondary Antibody Staining

The samples were subsequently incubated with Cy5-labeled secondary antibody (1:10,000 dilution) for 1 hour at room temperature. Following this, the slides were washed five times with 1X PBST.

#### Mounting and Imaging

A 5uL drop of DAPI solution was added to each slide, which was then covered with a coverslip. The samples were imaged using a fluorescence microscope.

#### Quantitation of IF2 foci

The cells were found through the DAPI channel and manually counted the number of cells per foci in the cy5 channel. The average fraction of cells having foci all three replicates was plotted on a bar graph and error bar is represented by standard error. The t-test (one-tailed distribution and two sample unequal variances) is used to estimate the statistical significance.

### Western Blot

For determining the spa tag in the strain *infB-SPA* strain^36^, both BL21(control) cells were grown in LB and *infB-SPA* strain cells were grown in LB-kanamycin media until it reached log phase. The cells were pelleted and resuspended in 250μl of 1x laemmli buffer for each 1.0 OD_600_ unit. The samples were boiled at 95C for 5 min then vortexed. 15ml of samples were loaded in a 10% SDS-PAGE gel and ran at 200V for 90 min. The PVDF membrane was incubated in 100% methanol for 15 s and rinsed in TBST buffer for 2 min. The transferring of the lysates from the SDS-PAGE gel to the PVDF membrane occurred by utilizing the BIO-RAD Trans-Blot Turbo Transfer system (1 amps, 188 2.5 volts, 30 minutes). Once the transfer was completed, the PVDF membrane was placed in a new container containing 5% non-fat dry milk in PBST buffer (blocking solution) at room temperature with gentle shaking. For primary antibody blotting, the membrane was submerged in (1:1000) dilution of the IF-2 antibody ((DYKDDDDK anti-flag tag) in the same blocking buffer and shaked gently for 1 hour at room temperature. After washing the membrane 3 times with the same PBST buffer for 10 min the membrane was probed with secondary antibody (1:10,000) anti-rabbit Cy5 secondary antibody (invitrogen) in blocking buffer. Secondary antibody incubation was done overnight with gentle shaking at 4°C temperature. The membrane was then washed 3 times, 10 min each, with 1x PBS buffer with gentle shaking. Following the wash, the membrane was then placed into an iBright imaging system, and the signal was detected under the Chemi-blot filter.

### Cell growth assays

*E. coli infB-SPA* strain and MG1655 cells were cultured in LB media at 37°C with shaking until reaching the log phase (OD600 = 0.5–0.6). The bacterial cultures were then diluted to an OD600 of 0.05 and further serially diluted 1:10 in LB media. 2.5 μL aliquots of each dilution were spotted on LB agar plates. The plates were incubated at 37°C overnight, and bacterial growth was assessed by observing colony formation the following day. Besides, comparison of both the liquid culture growth assay was also performed by measuring the optical density at 600 nm at regular intervals starting with OD= 0.05.

*E. coli* BL21infB::KanR pBAD-infB and *E. coli* BL21infB::KanR pBAD-infBΔNTD were cultured in LB media supplemented with ampicillin (60 μg/mL), kanamycin (25 μg/mL), and arabinose (0.1%) at 37°C with shaking until reaching the log phase (OD600 = 0.5–0.6). The bacterial cultures were then diluted to an OD600 of 0.05 and further serially diluted 1:10 in LB media. These dilutions were exposed to different cold stress conditions by incubating at 4°C, −20°C, and −80°C for 1 hour. After cold treatment, 2.5 μL aliquots of each dilution were spotted on LB agar plates containing ampicillin (60 μg/mL), kanamycin (25 μg/mL), and arabinose (0.1%). The plates were incubated at 37°C overnight, and bacterial growth was assessed by observing colony formation the following day.

### IF2 N-terminal IDR sequence analysis

#### Sequence retrieval and disorder prediction of all bacterial IF2 proteins

All available amino acid sequences of bacterial initiation factor 2 (IF2) from various bacterial species were downloaded in FASTA format from the UniProt database^37^. To assess intrinsic disorder, a Python script automated the submission of these sequences to the AIUPred server^38^, which provided disorder predictions for each residue in each sequence. The outputs were then processed again to calculate both the disorder length and the percentage of disordered residues for each IF2 sequence.

#### Phase separation prediction

The same set of IF2 sequences was analyzed using two phase separation predictors: deePhase^39^ and FuzDrop^40^. A Python script was used to obtain deePhase scores and FuzDrop scores for each sequence. Some sequences downloaded from the UniProt database contained “X” symbols representing unknown amino acids. Such sequences were not processed by the two phase separation predictors, as it led to an error.

#### Residue-specific disorder profiling of *E. coli* IF2

To obtain a detailed view of disorder across the IF2 protein, the *E. coli* IF2 sequence was downloaded separately from UniProt KB^37^. Residue-specific disorder scores were calculated for each amino acid position using AIUPred^38^. These scores were then plotted as a function of residue number to visualize disorder across the protein sequence.

#### Structural analysis

The *E. coli* IF2 AlphaFold-predicted structure was downloaded from the AlphaFold database^41^ to annotate the IF2 domain structure based on structural predictions. The AlphaFold PDB file of *E. coli* IF2 was made using VMD^42^ based on the AIUpred^38^ disorder score.

#### Psychrophile and thermophile sequence analysis

Select genomes from psychrophiles and thermophiles were identified by a literature search in pubmed, and their IF2 protein sequences were extracted. The protein sequence was used in Metapredict to acquire the intrinsically disordered regions.

## Results

### IF2 proteins contain a large N-terminal IDR

Proteins that can phase separate to form biomolecular condensates typically have large intrinsically disordered regions (IDRs). *E. coli* translation initiation factor 2 (IF2) was known to have a large IDR^5^, therefore we investigated the presence of the IDR across bacterial IF2 proteins. We downloaded 39197 bacterial IF2 protein sequences from uniprot^43^, and used AIUpred^38^ to estimate the presence of the IDR and the distribution of IDR lengths (Fig 1A). As a control, for bacterial proteins known to phase separate, we calculated IDR for *C. crescentus* RNase E, the core scaffold protein of BR-bodies^26^, biomolecular condensates that organize the mRNA decay machinery. As a negative control, we included *C. crescentus* PNPase, a protein that we confirmed cannot phase separate and which is a client protein of BR-bodies^44^. Across IF2 proteins, the range of IDR length and percentage of the protein which is disordered varied across a wide range, with a median of 320 amino acids and 35.98% disorder. This is slightly lower than the IDR length and percent disorder of *C. crescentus* RNase E, and dramatically higher than *C. crescentus* PNPase. To estimate whether IF2 proteins might phase separate, we utilized deePhase^39^ and FuzDrop^40^ software which predicts phase separation properties of proteins based upon their amino acid sequences. We observed that across both algorithms, IF2 proteins scored with a median of 0.72 on deePhase, and 0.42 on Fuzdrop, which is lower than *C. crescentus* RNase E (0.79 on deePhase and 0.69 on FuzDrop), but markedly higher than *C. crescentus* PNPase (0.24 on deePhase and 0.18 on FuzDrop). Overall, this suggested to us that IF2 proteins may have a conserved capability to phase separate. *E. coli* IF2 scored near the median across all IF2 proteins (320 amino acids IDR length, 36% disorder, 0.72 deePhase, 0.34 Fuzdrop), suggesting it was a good representative for further studies (Fig 1A,B). IF2 is known to have several domains, highlighted in 1B. While Domains IV-VI are essential and required for delivery of fMet-tRNA^fMet^ to the ribosome and GTP hydrolysis^45^, domains I-III comprising the IDR are non-essential and were found to be required for cell growth at 15°C^10^. While several structures of *E. coli* IF2 exist, the N-terminal IDR was notably absent, so we downloaded an alphafold3 structure^46^ to highlight the relative size of this IDR (Fig 1B). While the IDR in the alphafold3 structure contains very low confidence, we did notice some regions of predicted alpha-helical structure, in line with spectroscopy studies^5^. In summary, *E. coli* IF2 appears to have some of the hallmark features of phase separating proteins, making it a strong candidate for experimental follow up for its potential function as a biomolecular condensate.

**Figure 1.**
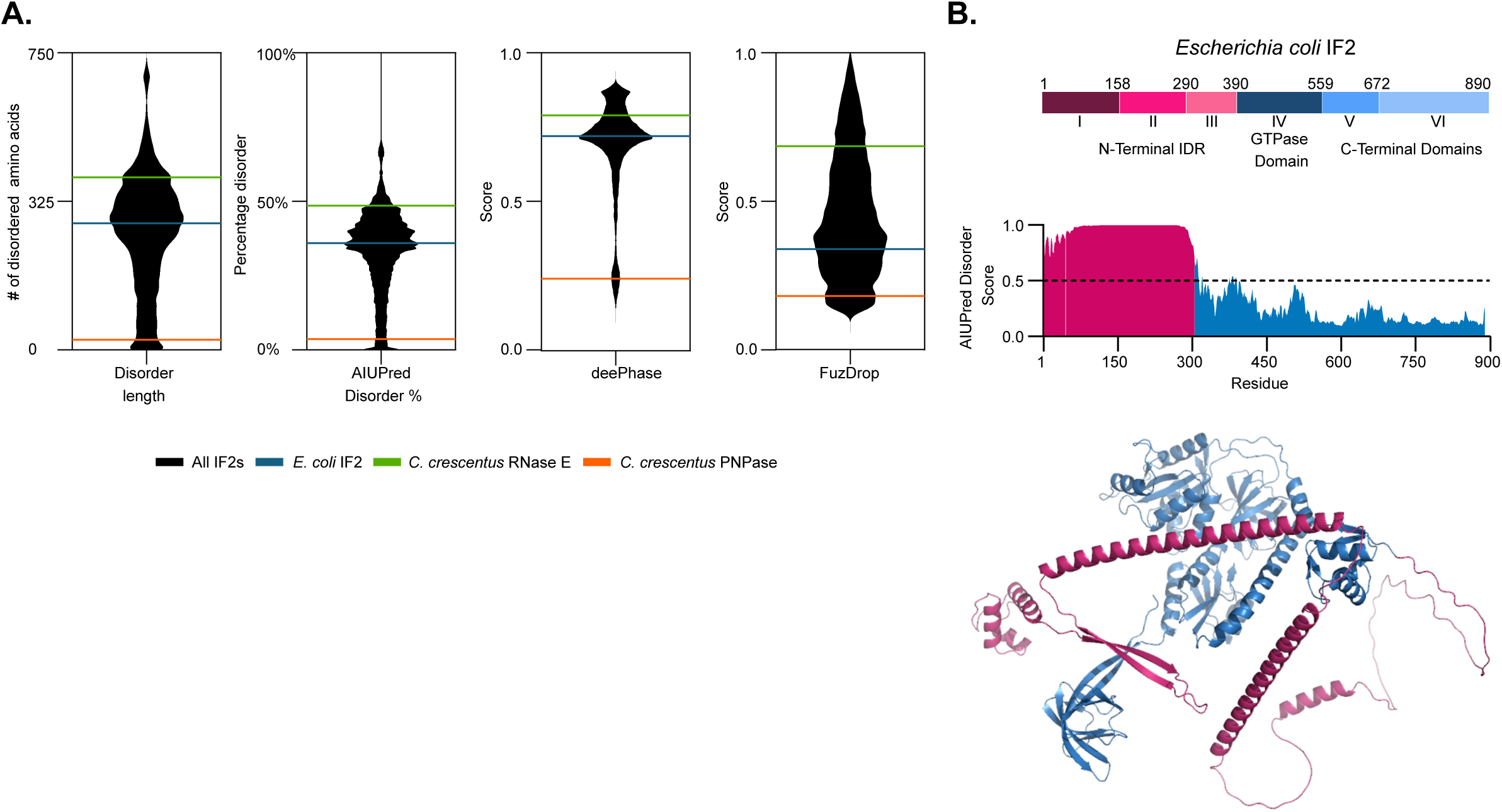
Bacterial IF2 has a large N-terminal IDR and is predicted to phase separate. **A)** A broad species-wide bioinformatics analysis of IF2 protein sequences across 39197 bacterial species. Disorder was predicted by AIUPred, and phase separation was predicted by deePhase and FuzDrop. The known phase separating protein *C. crescentus* RNase E, the core scaffold of BR-bodies (21), is the positive control while *C. crescentus* PNPase is the negative control. *E. coli* IF2 is indicated in Blue. **B)** Domain architecture of *E. coli* IF2. AlphaFold 2 structure shows *E. coli* IF2 contains a large intrinsically disordered region (highlighted in pink). High disorder in *E. coli* IF2 N-terminal region as predicted by AIUPred.

### IF2 phase separates with RNA

To experimentally determine whether *E. coli* IF2 can phase separate, we purified the α- isoform of IF2 (IF2 henceforth) and examined whether it could phase separate with *E. coli* total RNA (Fig 2A). IF2 was incubated at various concentrations and with various concentrations of RNA at room temperature, and we measured the turbidity and imaged the solution to examine whether IF2 phase separated (Fig S1). IF2 phase separated without RNA above a concentration of 5 µM, and we observed that the addition of total RNA reduced the critical concentration of IF2’s observed phase separation. To examine if IF2 droplets are reversible condensates, we subjected them to high salt (500mM KCl) to disrupt electrostatic interactions or RNase A treatment to degrade the RNA (Fig 2B,S2), which both appeared to dissolve IF2 droplets. These data suggest the observed IF2 droplets are biomolecular condensates which are in part assembled by electrostatic interactions and which RNA plays a role in driving phase separation. To further confirm the role that IF2 drives phase separation, we fused IF2 to a TEV cleavable solubilizing MBP tag ^47,48^. The MBP-IF2 fusion protein was unable to phase separate in the presence of total RNA, however, upon incubation with TEV protease we observed droplets, suggesting that IF2 is capable driving phase separation with RNA (Fig S3).

**Figure 2.**
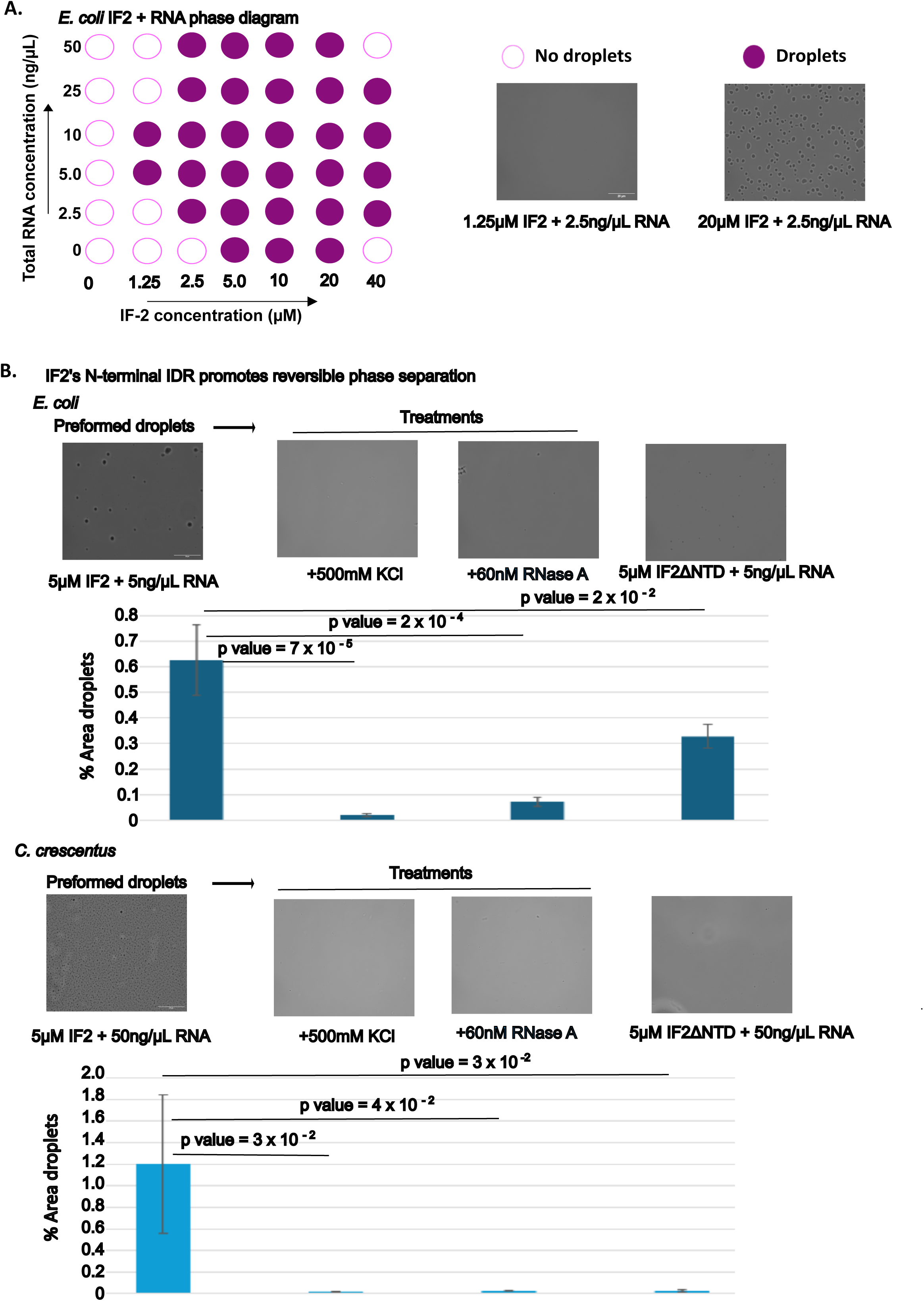
Phase Separation of IF-2 is stimulated by RNA and the N-terminal IDR. **A) Phase diagram of *E. coli* IF-2 and total RNA:** The two-component phase boundary for IF- 2_Ec and total RNA showing the result of concentration titrations. Turbidity measurements demonstrating phase separation of purified IF-2_Ec in the presence of total RNA. Phase-contrast microscope images of representative IF-2 RNA solutions showing that turbidity measures correspond to phase-separated condensates. **B) IF-2 Condensates are dissolved by 500mM KCl or RNase A treatment:** Purified IF-2 is incubated with 5ng total RNA in 20 mM Tris (pH 7.4), 75 mM KCl, 10 mM MgCl_2_, and 1 mM DTT for 30 minutes. IF-2 droplets were then treated with 500 mM KCl or 60nM RNase A for 30 minutes. Representative images from *E. coli* and *C. crescentus* shown. The value is the average of three independent replicates of > 50 droplets each, statistically significant is analyzed using t-test. The data were quantified in Image J software. **C) Enhanced Phase Separation by N-terminal IDR Regions:** Both *E. coli* and *Caulobacter crescentus* IF-2 proteins exhibit enhanced phase separation properties, driven by their intrinsically disordered regions. Representative images from *E. coli* and *C. crescentus* IF2 and IF-2 Δ N-terminal IDR are shown. The value is the average of three independent replicates of > 50 droplets each, statistically significant is analyzed using t-test. The data were quantified in Image J software.

To test the role of the N-terminal IDR on IF2’s observed phase separation, we purified a short form of the *E. coli* IF2 protein lacking the first 294 amino acids, called IF2ΔNTD. We observed IF2ΔNTD showed a notable reduction in IF2 droplets compared to the full length IF2 protein, suggesting the N-terminal IDR promotes IF2 phase separation. We also tested the β- isoform lacking the first 158 amino acids and found that the loss of this region leads to a robust loss in phase separation (Fig S5). To ensure that these IDR truncated variants were properly folded, we tested whether they were able to bind mant-GTP, a fluorescent analog of GTP. Indeed, all variants were able to bind mantGTP (Fig S4), suggesting the globular domain is properly folded. Altogether, this suggests that *E. coli* IF2 α-isoform can phase-separate with RNA, which is promoted by its N-terminal IDR.

Since IF2 proteins from various bacterial species also contain the IDR and are predicted to phase separate, we chose to explore whether IF2-RNA condensation is conserved across species. We chose to explore the α-proteobacterium *C. crescentus* IF2 which has a N-terminal IDR of 350 amino acids, and with 0.83 deePhase and 0.95 FuzDrop score respectively. We observed that *C. crescentus* IF2 protein phase separated with *C. crescentus* total RNA, and *C. crescentus* IF2 droplets were readily dissolved by high salt (500mM KCl) or RNase A treatment (Fig 2B). To investigate the role of *C. crescentus* IF2’s N-terminal IDR on phase separation, we also purified *C. crescentus* IF2ΔNTD protein and found that it almost completely lost all ability to phase separate (Fig 2B). Altogether, this suggests that IF2’s N-terminal IDR can promote phase separation across species, suggesting IF2-RNA condensation is likely a conserved function of the N-terminal IDR.

*T. thermophilus* IF2 was found to change from a compact state when bound to GDP, to an extended conformation when no nucleotide is bound^49^, suggesting nucleotide binding might influence phase separation. To test whether the nucleotide binding state impacts phase separation, we incubated IF2 with 1mM GTP, 1mM GDP, or no nucleotide and added total RNA (Fig S6). We observed that in the presence of GDP or GTP, IF2 showed a slight reduction in condensation in the presence of GTP, and no significant change in the presence of GDP (Fig S6). However, the condensates appeared to have irregular shapes and clump together in the presence of either GDP or GTP, suggesting potential maturation of the condensates into aggregates/tangle like structures. We chose to perform subsequent experiments in the absence of nucleotides where droplets remain spherical.

IF2 condensation occurs in the presence of total RNA, it was unclear which RNAs promote IF2 condensation. IF2 is known to directly bind to ribosomes and initiator tRNA, so we investigated the role of these RNAs on IF2 phase separation using purified *E. coli* molecules. IF2 incubated with only 70S ribosomes or only tRNA^fMet^ did not phase separate in our hands (data not shown). Interestingly, we observed that IF2-total-RNA droplets had reduced phase separation if we included 70S ribosomes or tRNA^fMet^ together with IF2 (Fig 2A). To examine whether 70S ribosomes or tRNA^fMet^ could dissolve IF2-total-RNA droplets, we pre-formed IF2- total-RNA droplets and then added 70S ribosomes or tRNA^fMet^. We observed that 70S ribosomes and tRNA^fMet^ could both dissolve IF2-total-RNA droplets (Fig 3A), suggesting that both IF2 binding partners reduce IF2-total-RNA phase separation.

**Figure 3.**
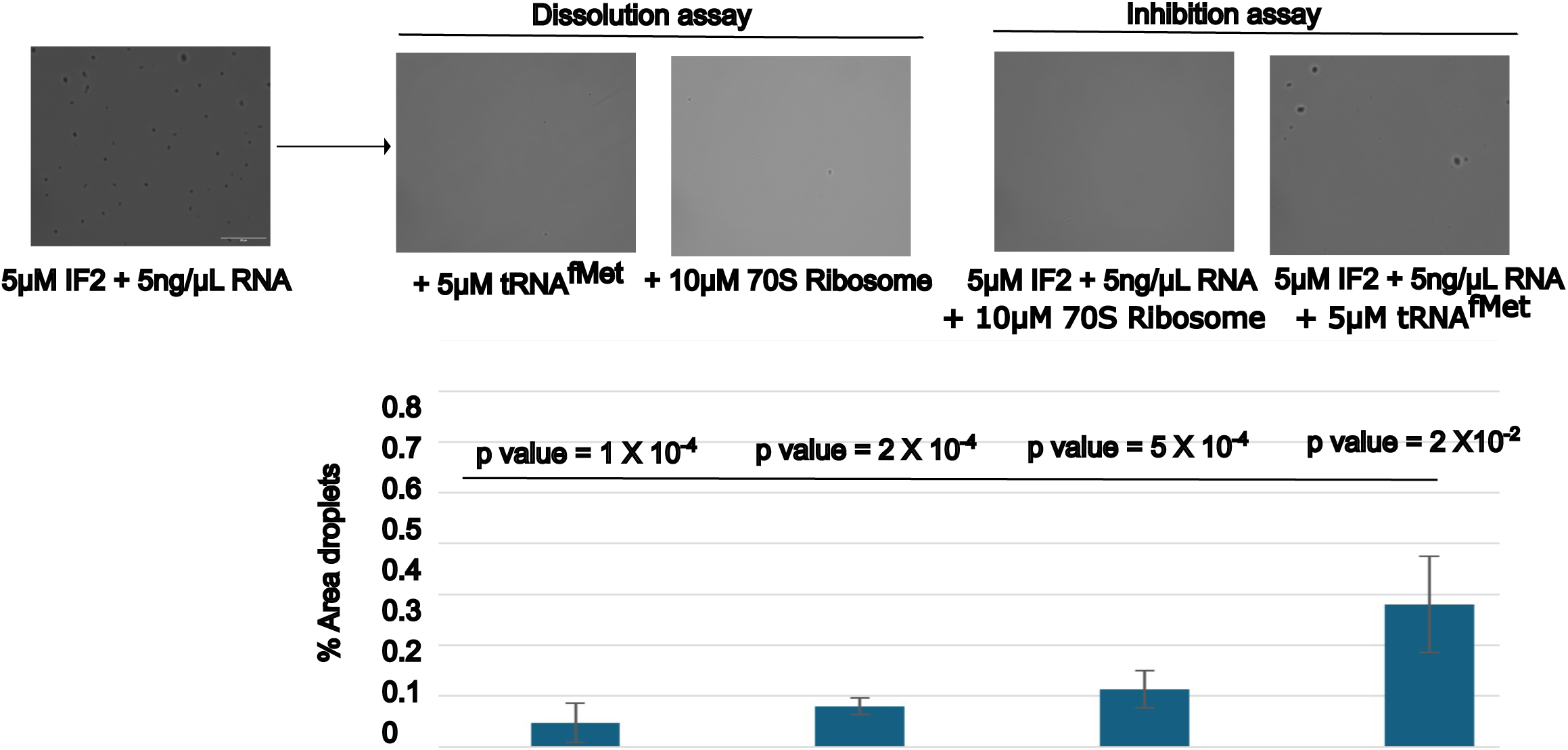
70S Ribosomes and tRNA^fMet^ reduce IF-2 phase separation. **A) IF-2 condensation is inhibited by ribosome and tRNA^fMet^:** Purified 5 μM IF-2 incubated with 5 ng/μL total RNA in 20 mM Tris (pH 7.4), 75 mM KCl, 10 mM MgCl_2_, and 1 mM DTT. Additional reactions were incubated with 10 μM 70S ribosomes or 5 μM tRNA^fMet^. Top: bar graph quantifying the percentage of image area covered in IF-2 RNA droplets using imageJ. The value is the average of three independent replicates of > 50 droplets each, statistically significant is analyzed using t-test (one-tailed distribution and unequal variance). Below: Representative images of IF-2 RNA droplets with different added components. **B) The IF-2 RNA condensates can be dissolved by the ribosome and tRNA^fMet^**: 5μM IF-2 condensate is formed with 5ng/μL total RNA in 20 mM Tris (pH 7.4), 75 mM KCl, 10 mM MgCl_2_, and 1 mM DTT for 30 minutes followed by the addition of 10μM of 70S ribosomes or 5μM of tRNA^fMet^ reduces the phase separation.

### N-terminal IDR promotes cold-induced phase separation and fitness during cold-shock

As *E. coli* IF2 lacking the N-terminal IDR was found to be unable to grow at lower temperatures^10^, we investigated the role of temperature on IF2-total-RNA phase separation (Fig 4). We incubated *E. coli* IF2 and total RNA at various temperatures, from the *E. coli* growth temperature (37°C), room temperature (25°C), and at 4°C (Fig 4A). Compared to room temperature, we observed that IF2 phase separation was reduced at 37°C and was strongly stimulated at 4°C. To investigate the role of the N-terminal IDR on cold-induced phase separation, we compared full length IF2 to IF2ΔNTD at 4°C (Fig 4B). Compared to full length IF2, we saw that IF2ΔNTD-total-RNA phase separation was strongly reduced, suggesting that the N-terminal IDR is required for phase separation at 4°C. To investigate if IF2-total-RNA droplets are reversible condensates at 4°C, we raised the temperature from 4°C to room temperature and observed that the IF2 phase separation was reduced after raising the temperature to 25°C (Fig 4B). Taken together, this suggests that IF2-total-RNA droplets are stimulated by the N-terminal IDR at low temperatures.

**Figure 4.**
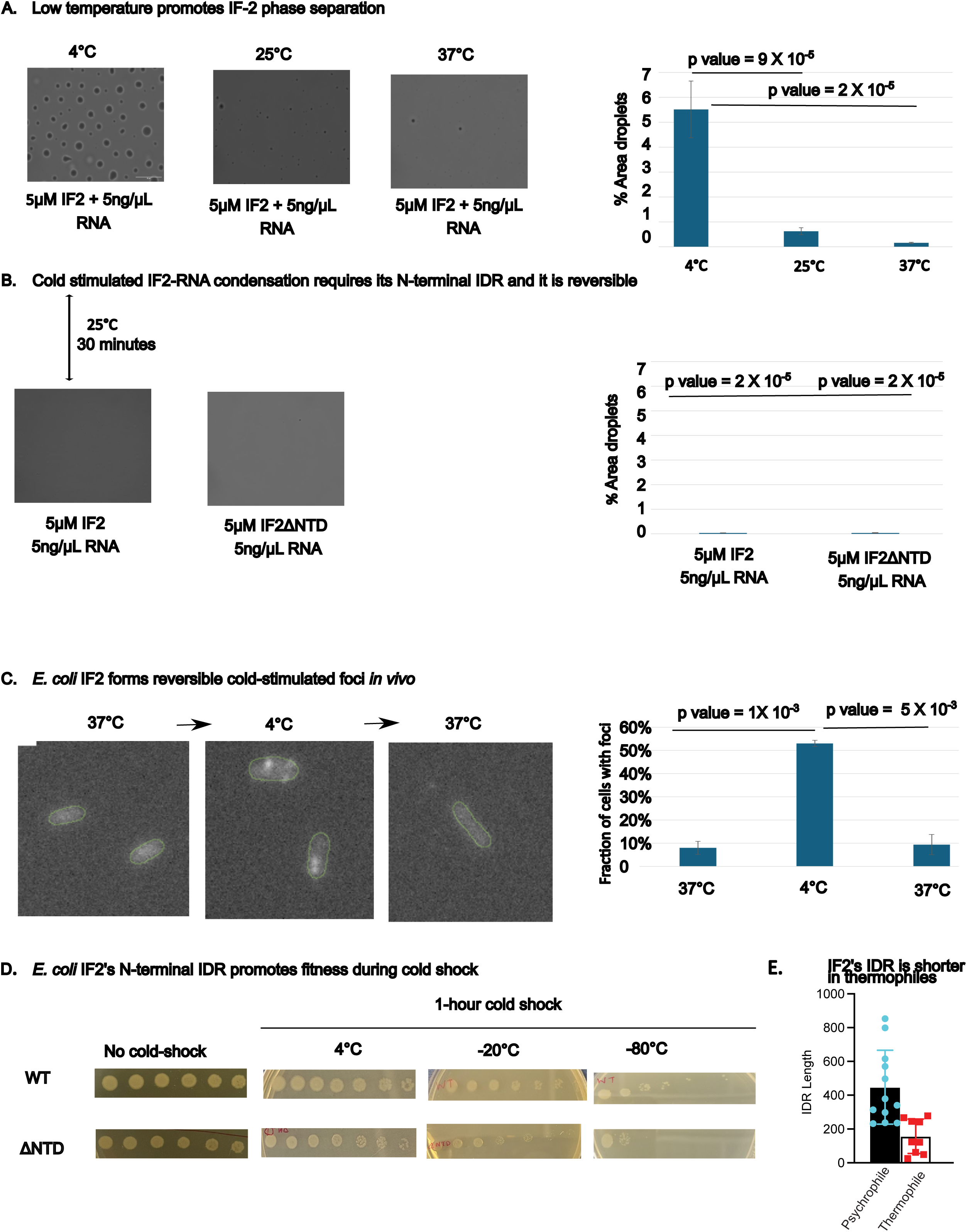
The N-terminal IDR promotes cold-driven phase separation and fitness during cold-shock. **A) Phase contrast imaging of purified IF-2 RNA condensates at various temperatures:** The purified IF-2 is incubated with 5ng/uL total RNA in 20 mM Tris (pH 7.4), 75 mM KCl, 10 mM MgCl2, and 1 mM DTT buffer at 4°C, 25°C and 37°C. Phase contrast images showing that cold temperatures (4°C) induce more robust phase separation of IF-2 with total RNA compared to room temperature (RT) and 37°C. The value is the average of three independent replicates of > 50 droplets each, statistically significant is analyzed using a 1-sided t-test with uneven variance. The data were quantified in Image J software. **B) N-terminal IDR is required for cold-induced phase separation:** The purified IF-2 and ΔNTD IF-2 is incubated with 5 ng/μL total RNA in 20 mM Tris (pH 7.4), 75 mM KCl, 10 mM MgCl2, and 1 mM DTT buffer at 4°C. The value is the average of three independent replicates of > 50 droplets each, statistically significant is analyzed using a 1-sided t-test with uneven variance. The data were quantified in Image J software. **C) IF-2 foci are reversibly induced in cold temperature *in vivo*:** Immunofluorescence imaging performed on *E. coli infB::FLAG* (ref) cells fixed in log growth at 37°C, after 1 hour at 4°C, or after returning the cells to 37°C for 1 hour. Scale bar is 1 µm. Green outline of cell was generated using microbeJ (ref). **D) The N-terminal IDR promotes fitness under cold shock:** *E. coli* cells were grown to the log phase (OD=0.5) in Luria broth at 37°C in a shaker-incubator and subjected to a 1-hour cold shock at 4°C, −20°C, or −80°C. After cold shock, we plated 5 μL of serial dilutions on LB kanamycin, ampicillin arabinose plates. The plates were then incubated at 37°C overnight to allow for growth before imaging. **E) The N-terminal IDR length is reduced in thermophiles.** Psychrophilic and thermophilic bacteria were identified, and the sequence lengths were plotted (Species listed in Fig S10).

To investigate whether IF2 condensates occur *in vivo*, we performed immunofluorescence using an *infB-SPA* strain^36^. We compared the growth rates of *infB-SPA* to wild type *E. coli*, and observed similar growth (Fig S7), suggesting that the FLAG tag does not impact IF2 function. We therefore grew *infB-SPA* cells at the standard growth temperature (37°C), or after 1 hour of cold-shock at 4°C. The cells were then fixed with formaldehyde, permeabilized, and probed with anti-FLAG primary antibody, followed by a Cy5-labeled secondary antibody (Fig 4C). First, we confirmed that the IF2 antibody was selective, and we observed no staining in the absence of the SPA tag (Fig S8), and that IF2-SPA α-isoform was the major band observed by western blot (Fig S9). At the standard growth temperature, we observed that IF2 was diffuse across the cytoplasm, however, when cells were cold-shocked for 1-hour we observed that IF2 localized into distinct puncta (Fig 4C). To examine if the IF2 foci observed at 4°C were reversible condensates, we recovered the cells at 37°C for 1 hour and performed fixation. After recovery from 4°C, we observed that IF2 foci were lost (Fig 4C), suggesting that IF2 foci are reversible *in vivo*. To investigate whether IF2’s N-terminal IDR promotes fitness during cold shock, we grew cultures of *E. coli* expressing a single copy of either full length IF2 or IF2ΔNTD from a plasmid^10^ at 37°C, and subjected the cells to 1-hour cold shocks at 4°C, - 20°C, or −80°C (Fig 4D). Upon growth at 37°C, we observed no difference between the two strains. Upon cold-shock, we noticed that the IF2ΔNTD strain showed a significant reduction in fitness compared to the IF2 strain that was small at 4°C but grew bigger at −20°C and was large at −80°C. As the severity of cold-shock increased, we observed a decrease in CFU’s in the IF2ΔNTD strain relative to IF2, suggesting that the N-terminal IDR promotes fitness to *E. coli* cells during cold shock. In the same genetic system, it was previously established that the IF2ΔNTD was unable to grow at 15°C ^10^. As IDR length varied significantly across species, we explored whether the N-terminal IDR length varied across different species with distinct optimal growth temperatures. Therefore, we collected a set of IF2s derived from psychrophile and thermophile bacteria and calculated their IDR lengths (Fig 4E). Here we observed that psychrophiles tended to have longer IDRs, while thermophiles have much shorter IDRs, in-line with our experimentally derived role in promoting fitness at lower temperatures in *E. coli*.

## Discussion

The N-terminal IDR of bacterial IF2 is capable of driving phase separation with cellular RNA into a biomolecular condensate, suggesting a conserved molecular function of the IDR. Interestingly, IF2 IDRs tend to share low sequence identity, with *E. coli* and *C. crescentus* IF2s sharing only 40.8% amino acid identity, yet they are both capable of phase separation. Across bacteria, most IF2 sequences are predicted to have the capacity to phase separate by deePhase^39^ or fuzdrop^40^ algorithms, so it will be important to investigate the molecular and functional properties of IF2 condensates across species.

Bacterial IF2 condensates share many similarities with cold-induced eukaryotic stress granules^25^, suggesting they may play a similar functional role across domains. Unlike constitutive bacterial condensates like BR-bodies^26^, IF2 condensates are not present at normal growth temperature, but are induced by a shift to cold temperature^25^. Based upon the genetic system established in *E. coli* cells^10^, we have found that IF2 has enhanced phase separation in the cold, and that IF2 condensates appear to protect *E. coli* cells during adaptation to the cold (Fig. 4D), where the N-terminal IDR was found to be required for growth at cold temperatures^10^. Interestingly, phosphorylation of eIF2α by the integrated stress response triggering reduced translation is important for eukaryotic stress granule formation^25,50^, placing IF2 as a key player in bacteria and eukaryote cold-induced condensates. Interestingly, bacterial IF2 is encoded by a single protein, while in eukaryotes it is encoded by three separate protein subunits (eIF2α, eIF2β, and eIF2γ). While eukaryotic stress granules are induced by the cold, they can also be induced by other stresses including heat shock, oxidative stress, viral infection, osmotic stress, and UV irradiation^23^. Therefore, an important goal will be to determine whether bacterial IF2 condensates are induced by similar stresses and whether they promote fitness across stresses. It is well established that cold-shock in *E. coli* leads to a strong reduction in translation^51,52^, and an important goal moving forward will be to determine whether IF2 condensation is involved in the translation shutdown during adaptation to the cold. In line with their role in the cold, we observed that IF2’s IDR length is significantly reduced in thermophiles (Fig 4E). While the RNA composition of cold-induced IF2 condensates is not yet known, an important goal will be to identify which cellular RNAs and proteins are contained in IF2 condensates. Interestingly, lamotrigine, a novel antibiotic candidate, has been found to already bind to the N-terminal IDR and selectively inhibit *E. coli* growth in the cold^11^. This suggests IF2 condensates present a promising target for the development of next-generation antibiotics.

## Resource availability

All strains and plasmids generated in this manuscript are available upon request to the lead contact.

## Acknowledgements

NIH NIGMS grant R35GM124733 and IU startup funds to JMS. NIH Grant R01GM136863 to WSC. Work reported in this publication was supported by the National Institutes of Health Common Fund and Office of Scientific Workforce Diversity under three linked awards RL5GM118981, TL4GM118983, and 1UL1GM118982 administered by the National Institute of General Medical Sciences for NRN. Dr. Tamara Hendrickson and the WSU department of chemistry for lab space and equipment after the Schrader lab was destroyed in a fire. Dr. Hannah Hariri for use of their microscope. Dr. Claudio Gualerzi for providing *E. coli* Strains.

## Author contributions

JMS and WSC acquired funding, designed study, and managed the research team. AG contributed to study design and performed *in vitro* and *in vivo* experiments. KSM and KD performed IF2 protein sequence analysis. VN performed MANT-GTP binding experiments. NN cloned the *C. crescentus* IF2 expression plasmid. JMS, WSC, AG, KSM, KGD, and VN analyzed data. JMS and AG wrote the manuscript.

## References

1. Milon, P., and Rodnina, M.V. (2012). Kinetic control of translation initiation in bacteria. Crit. Rev. Biochem. Mol. Biol. 47, 334–348. 10.3109/10409238.2012.678284.

2. Rodnina, M.V. (2018). Translation in Prokaryotes. Cold Spring Harb. Perspect. Biol. 10, a032664. 10.1101/cshperspect.a032664.

3. Gualerzi, C.O., and Pon, C.L. (2015). Initiation of mRNA translation in bacteria: structural and dynamic aspects. Cell. Mol. Life Sci. CMLS 72, 4341–4367. 10.1007/s00018-015-2010-3.

4. Laalami, S., Sacerdot, C., Vachon, G., Mortensen, K., Sperling-Petersen, H.U., Cenatiempo, Y., and Grunberg-Manago, M. (1991). Structural and functional domains of *E coli* initiation factor IF2. Biochimie 73, 1557–1566. 10.1016/0300-9084(91)90191-3.

5. Laursen, B.S., Kjærgaard, A.C., Mortensen, K.K., Hoffman, D.W., and Sperling-Petersen, H.U. (2004). The N-terminal domain (IF2N) of bacterial translation initiation factor IF2 is connected to the conserved C-terminal domains by a flexible linker. Protein Sci. Publ. Protein Soc. 13, 230–239. 10.1110/ps.03337604.

6. Simonetti, A., Marzi, S., Billas, I.M.L., Tsai, A., Fabbretti, A., Myasnikov, A.G., Roblin, P., Vaiana, A.C., Hazemann, I., Eiler, D., et al. (2013). Involvement of protein IF2 N domain in ribosomal subunit joining revealed from architecture and function of the full-length initiation factor. Proc. Natl. Acad. Sci. U. S. A. 110, 15656–15661. 10.1073/pnas.1309578110.

7. Laursen, B.S., Mortensen, K.K., Sperling-Petersen, H.U., and Hoffman, D.W. (2003). A conserved structural motif at the N terminus of bacterial translation initiation factor IF2. J. Biol. Chem. 278, 16320–16328. 10.1074/jbc.M212960200.

8. Plumbridge, J.A., Deville, F., Sacerdot, C., Petersen, H.U., Cenatiempo, Y., Cozzone, A., Grunberg-Manago, M., and Hershey, J.W. (1985). Two translational initiation sites in the infB gene are used to express initiation factor IF2 alpha and IF2 beta in Escherichia coli. EMBO J. 4, 223–229. 10.1002/j.1460-2075.1985.tb02339.x.

9. Cenatiempo, Y., Deville, F., Dondon, J., Grunberg-Manago, M., Sacerdot, C., Hershey, J.W., Hansen, H.F., Petersen, H.U., Clark, B.F., and Kjeldgaard, M. (1987). The protein synthesis initiation factor 2 G-domain. Study of a functionally active C-terminal 65-kilodalton fragment of IF2 from Escherichia coli. Biochemistry 26, 5070–5076. 10.1021/bi00390a028.

10. Brandi, A., Piersimoni, L., Feto, N.A., Spurio, R., Alix, J.-H., Schmidt, F., and Gualerzi, C.O. (2019). Translation initiation factor IF2 contributes to ribosome assembly and maturation during cold adaptation. Nucleic Acids Res. 47, 4652–4662. 10.1093/nar/gkz188.

11. Stokes, J.M., Davis, J.H., Mangat, C.S., Williamson, J.R., and Brown, E.D. (2014). Discovery of a small molecule that inhibits bacterial ribosome biogenesis. eLife 3, e03574. 10.7554/eLife.03574.

12. Madison, K.E., Abdelmeguid, M.R., Jones-Foster, E.N., and Nakai, H. (2012). A New Role for Translation Initiation Factor 2 in Maintaining Genome Integrity. PLoS Genet. 8, e1002648. 10.1371/journal.pgen.1002648.

13. Mallikarjun, J., and Gowrishankar, J. Essential Role for an Isoform of Escherichia coli Translation Initiation Factor IF2 in Repair of Two-Ended DNA Double-Strand Breaks. J. Bacteriol. 204, e00571–21. 10.1128/jb.00571-21.

14. Banani, S.F., Lee, H.O., Hyman, A.A., and Rosen, M.K. (2017). Biomolecular condensates: organizers of cellular biochemistry. Nat. Rev. Mol. Cell Biol. 18, 285–298. 10.1038/nrm.2017.7.

15. Lafontaine, D.L.J., Riback, J.A., Bascetin, R., and Brangwynne, C.P. (2021). The nucleolus as a multiphase liquid condensate. Nat. Rev. Mol. Cell Biol. 22, 165–182. 10.1038/s41580-020-0272-6.

16. Brangwynne, C.P., Eckmann, C.R., Courson, D.S., Rybarska, A., Hoege, C., Gharakhani, J., Jülicher, F., and Hyman, A.A. (2009). Germline P granules are liquid droplets that localize by controlled dissolution/condensation. Science 324, 1729–1732. 10.1126/science.1172046.

17. Blake, L.A., Watkins, L., Liu, Y., Inoue, T., and Wu, B. (2024). A rapid inducible RNA decay system reveals fast mRNA decay in P-bodies. Nat. Commun. 15, 2720. 10.1038/s41467-024-46943-z.

18. Currie, S.L., Xing, W., Muhlrad, D., Decker, C.J., Parker, R., and Rosen, M.K. (2023). Quantitative reconstitution of yeast RNA processing bodies. Proc. Natl. Acad. Sci. U. S. A. 120, e2214064120. 10.1073/pnas.2214064120.

19. Iserman, C., Desroches Altamirano, C., Jegers, C., Friedrich, U., Zarin, T., Fritsch, A.W., Mittasch, M., Domingues, A., Hersemann, L., Jahnel, M., et al. (2020). Condensation of Ded1p Promotes a Translational Switch from Housekeeping to Stress Protein Production. Cell 181, 818–831.e19. 10.1016/j.cell.2020.04.009.

20. Riback, J.A., Katanski, C.D., Kear-Scott, J.L., Pilipenko, E.V., Rojek, A.E., Sosnick, T.R., and Drummond, D.A. (2017). Stress-Triggered Phase Separation Is an Adaptive, Evolutionarily Tuned Response. Cell 168, 1028–1040 e19. 10.1016/j.cell.2017.02.027.

21. Jain, S., Wheeler, J.R., Walters, R.W., Agrawal, A., Barsic, A., and Parker, R. (2016). ATPase-Modulated Stress Granules Contain a Diverse Proteome and Substructure. Cell 164, 487–498. 10.1016/j.cell.2015.12.038.

22. Khong, A., Matheny, T., Jain, S., Mitchell, S.F., Wheeler, J.R., and Parker, R. (2017). The Stress Granule Transcriptome Reveals Principles of mRNA Accumulation in Stress Granules. Mol. Cell 68, 808–820 e5. 10.1016/j.molcel.2017.10.015.

23. Hofmann, S., Kedersha, N., Anderson, P., and Ivanov, P. (2021). Molecular mechanisms of stress granule assembly and disassembly. Biochim. Biophys. Acta Mol. Cell Res. 1868, 118876. 10.1016/j.bbamcr.2020.118876.

24. Protter, D.S.W., and Parker, R. (2016). Principles and Properties of Stress Granules. Trends Cell Biol 26, 668–679. 10.1016/j.tcb.2016.05.004.

25. Hofmann, S., Cherkasova, V., Bankhead, P., Bukau, B., and Stoecklin, G. (2012). Translation suppression promotes stress granule formation and cell survival in response to cold shock. Mol. Biol. Cell 23, 3786–3800. 10.1091/mbc.E12-04-0296.

26. Al-Husini, N., Tomares, D.T., Bitar, O., Childers, W.S., and Schrader, J.M. (2018). α-Proteobacterial RNA Degradosomes Assemble Liquid-Liquid Phase-Separated RNP Bodies. Mol. Cell 71, 1027–1039.e14. 10.1016/j.molcel.2018.08.003.

27. Azaldegui, C.A., Vecchiarelli, A.G., and Biteen, J.S. (2021). The emergence of phase separation as an organizing principle in bacteria. Biophys. J. 120, 1123–1138. 10.1016/j.bpj.2020.09.023.

28. Nandana, V., and Schrader, J.M. (2021). Roles of liquid–liquid phase separation in bacterial RNA metabolism. Curr. Opin. Microbiol. 61, 91–98. 10.1016/j.mib.2021.03.005.

29. Ladouceur, A.-M., Parmar, B.S., Biedzinski, S., Wall, J., Tope, S.G., Cohn, D., Kim, A., Soubry, N., Reyes-Lamothe, R., and Weber, S.C. (2020). Clusters of bacterial RNA polymerase are biomolecular condensates that assemble through liquid–liquid phase separation. Proc. Natl. Acad. Sci. 117, 18540–18549. 10.1073/pnas.2005019117.

30. Al-Husini, N., Tomares, D.T., Pfaffenberger, Z.J., Muthunayake, N.S., Samad, M.A., Zuo, T., Bitar, O., Aretakis, J.R., Bharmal, M.-H.M., Gega, A., et al. (2020). BR-bodies provide selectively permeable condensates that stimulate mRNA decay and prevent release of decay intermediates. Mol. Cell 78, 670–682.e8. 10.1016/j.molcel.2020.04.001.

31. Goldberger, O., Szoke, T., Nussbaum-Shochat, A., and Amster-Choder, O. (2022). Heterotypic phase separation of Hfq is linked to its roles as an RNA chaperone. Cell Rep. 41, 111881. 10.1016/j.celrep.2022.111881.

32. Beaufay, F., Amemiya, H.M., Guan, J., Basalla, J., Meinen, B.A., Chen, Z., Mitra, R., Bardwell, J.C.A., Biteen, J.S., Vecchiarelli, A.G., et al. (2021). Polyphosphate drives bacterial heterochromatin formation. Sci. Adv. 7, eabk0233. 10.1126/sciadv.abk0233.

33. Mittag, T., and Pappu, R.V. (2022). A conceptual framework for understanding phase separation and addressing open questions and challenges. Mol. Cell 82, 2201–2214. 10.1016/j.molcel.2022.05.018.

34. Boeynaems, S., Alberti, S., Fawzi, N.L., Mittag, T., Polymenidou, M., Rousseau, F., Schymkowitz, J., Shorter, J., Wolozin, B., Van Den Bosch, L., et al. (2018). Protein Phase Separation: A New Phase in Cell Biology. Trends Cell Biol 28, 420–435. 10.1016/j.tcb.2018.02.004.

35. Butland, G., Peregrín-Alvarez, J.M., Li, J., Yang, W., Yang, X., Canadien, V., Starostine, A., Richards, D., Beattie, B., Krogan, N., et al. (2005). Interaction network containing conserved and essential protein complexes in Escherichia coli. Nature 433, 531–537. 10.1038/nature03239.

36. Butland, G., Peregrin-Alvarez, J.M., Li, J., Yang, W., Yang, X., Canadien, V., Starostine, A., Richards, D., Beattie, B., Krogan, N., et al. (2005). Interaction network containing conserved and essential protein complexes in Escherichia coli. Nature 433, 531–537. 10.1038/nature03239.

37. The UniProt Consortium (2023). UniProt: the Universal Protein Knowledgebase in 2023. Nucleic Acids Res. 51, D523–D531. 10.1093/nar/gkac1052.

38. Erdős, G., and Dosztányi, Z. (2024). AIUPred: combining energy estimation with deep learning for the enhanced prediction of protein disorder. Nucleic Acids Res. 52, W176–W181. 10.1093/nar/gkae385.

39. Saar, K.L., Morgunov, A.S., Qi, R., Arter, W.E., Krainer, G., Lee, A.A., and Knowles, T.P.J. (2021). Learning the molecular grammar of protein condensates from sequence determinants and embeddings. Proc. Natl. Acad. Sci. U. S. A. 118, e2019053118. 10.1073/pnas.2019053118.

40. Hatos, A., Tosatto, S.C.E., Vendruscolo, M., and Fuxreiter, M. (2022). FuzDrop on AlphaFold: visualizing the sequence-dependent propensity of liquid-liquid phase separation and aggregation of proteins. Nucleic Acids Res. 50, W337–W344. 10.1093/nar/gkac386.

41. Varadi, M., Anyango, S., Deshpande, M., Nair, S., Natassia, C., Yordanova, G., Yuan, D., Stroe, O., Wood, G., Laydon, A., et al. (2022). AlphaFold Protein Structure Database: massively expanding the structural coverage of protein-sequence space with high-accuracy models. Nucleic Acids Res. 50, D439–D444. 10.1093/nar/gkab1061.

42. Humphrey, W., Dalke, A., and Schulten, K. (1996). VMD: visual molecular dynamics. J. Mol. Graph. 14, 33–38, 27–28. 10.1016/0263-7855(96)00018-5.

43. UniProt: a hub for protein information (2015). Nucleic Acids Res 43, D204–12. 10.1093/nar/gku989.

44. Nandana, V., Rathnayaka-Mudiyanselage, I.W., Muthunayake, N.S., Hatami, A., Mousseau, C.B., Ortiz-Rodríguez, L.A., Vaishnav, J., Collins, M., Gega, A., Mallikaarachchi, K.S., et al. (2023). The BR- body proteome contains a complex network of protein-protein and protein-RNA interactions. Cell Rep. 42, 113229. 10.1016/j.celrep.2023.113229.

45. Spurio, R., Brandi, L., Caserta, E., Pon, C.L., Gualerzi, C.O., Misselwitz, R., Krafft, C., Welfle, K., and Welfle, H. (2000). The C-terminal subdomain (IF2 C-2) contains the entire fMet-tRNA binding site of initiation factor IF2. J. Biol. Chem. 275, 2447–2454. 10.1074/jbc.275.4.2447.

46. Jumper, J., Evans, R., Pritzel, A., Green, T., Figurnov, M., Ronneberger, O., Tunyasuvunakool, K., Bates, R., Žídek, A., Potapenko, A., et al. (2021). Highly accurate protein structure prediction with AlphaFold. Nature 596, 583–589. 10.1038/s41586-021-03819-2.

47. Brumbaugh-Reed, E.H., Gao, Y., Aoki, K., and Toettcher, J.E. (2024). Rapid and reversible dissolution of biomolecular condensates using light-controlled recruitment of a solubility tag. Nat. Commun. 15, 6717. 10.1038/s41467-024-50858-0.

48. Lin, Y., Protter, D.S.W., Rosen, M.K., and Parker, R. (2015). Formation and Maturation of Phase-Separated Liquid Droplets by RNA-Binding Proteins. Mol. Cell 60, 208–219. 10.1016/j.molcel.2015.08.018.

49. Eiler, D., Lin, J., Simonetti, A., Klaholz, B.P., and Steitz, T.A. (2013). Initiation factor 2 crystal structure reveals a different domain organization from eukaryotic initiation factor 5B and mechanism among translational GTPases. Proc. Natl. Acad. Sci. 110, 15662–15667. 10.1073/pnas.1309360110.

50. English, A.M., Green, K.M., and Moon, S.L. (2022). A (dis)integrated stress response: Genetic diseases of eIF2α regulators. Wiley Interdiscip. Rev. RNA 13, e1689. 10.1002/wrna.1689.

51. Zhang, Y., and Gross, C.A. (2021). Cold Shock Response in Bacteria. Annu. Rev. Genet. 55, 377–400. 10.1146/annurev-genet-071819-031654.

52. Zhang, Y., Burkhardt, D.H., Rouskin, S., Li, G.-W., Weissman, J.S., and Gross, C.A. (2018). A Stress Response that Monitors and Regulates mRNA Structure Is Central to Cold Shock Adaptation. Mol. Cell 70, 274–286.e7. 10.1016/j.molcel.2018.02.035.

