## Supplemental information for "Bacterial IF2’s N-terminal IDR drives cold-induced phase separation and promotes fitness during cold stress"

### Supplementary Figure Legends

**Fig S1.** Bar graphs show turbidity increases with increasing IF-2 concentration, reaching a plateau at the basal IF-2 concentration (20 µM) in the absence of RNA. In the presence of varying RNA concentrations, turbidity initially increases but decreases at higher RNA concentrations, indicating a concentration-dependent modulation of phase separation by RNA.

**Fig S2.** In the presence of 5 µM IF-2 and 5 ng/µL total RNA, an increase in turbidity was observed. However, the addition of excess KCl or RNase A resulted in a decrease in turbidity, indicating that the turbidity induced by the IF-2 protein is reversible under these conditions.

**Fig S3.** 20 µM IF-2 tagged MBP protein was incubated in a buffer containing 20 mM Tris (pH 7.4), 75 mM KCl, 10 mM $MgCl_2$, and 1 mM DTT for 2.5 hours at room temperature. Under these conditions, the protein did not undergo phase separation. However, upon the addition of TEV protease at a 1:15 molar ratio and incubation for 2.5 hours, the IF-2 protein exhibited clear phase separation. This suggests that the cleavage of the MBP tag by TEV protease promotes the phase separation of IF-2.

**Fig S4.** 125 µM MANT-GTP was incubated with 1 µM IF2, 1 µM IF2ΔNTD, 1 µM IF2β, or 1 µM Aconitase and measured in a fluorescence plate reader. Fluorescence enhancement at 450 nm was calculated as a ratio of the MANT-GTP fluorescence in the presence of each protein relative relative to the MANT-GTP fluorescence intensity. As a control, proteins samples were also incubated with 3mM GTP as an unlabeled chase. Protein samples lacking MANT-GTP were run as negative controls. Error bars represent standard deviation from three independent replicate measurements. * indicates $p<0.05$ from 1 sided t-test with uneven variance.

**Fig S5.** Purified 5 µM IF-2 β isoforms was incubated with 5 ng/µL total RNA in a buffer containing 20 mM Tris (pH 7.4), 75 mM KCl, 10 mM MgCl2, and 1 mM DTT at room temperature. The bar graph represents the percentage of image area covered by IF-2 RNA droplets, quantified using ImageJ. Values shown are the averages of three independent replicates, each consisting of >50 droplets. Statistically significant differences between full length and β isoforms were determined using a one-tailed t-test for samples with unequal variance.

**Fig S6.** Purified 5 µM IF-2 was incubated with 5 ng/µL total RNA and 1 mM GTP or GDP in a buffer containing 20 mM Tris (pH 7.4), 75 mM KCl, 10 mM MgCl2, and 1 mM DTT at room temperature. The presence of nucleotides led to a reduction in droplet formation or the appearance of aggregates. The bar graph represents the percentage of image area covered by IF-2 RNA droplets, quantified using ImageJ. Values shown are the averages of three independent replicates, each consisting of >50 droplets. Statistically significant differences between conditions with and without nucleotides were determined using a one-tailed t-test for samples with unequal variance.

**Fig S7.** Comparison of the growth assay between the infB-SPA strain and the wild-type MG1655 strain. Cells were grown to log phase ($OD_{600}$ = 0.5), serially diluted in a 1:10 ratio, and spotted onto plates. The plates were incubated overnight at 37°C. No growth difference was observed between the tagged infB-SPA strain and the MG1655 wild-type strain. The liquid growth curve assay also shows that there is no significant difference between them and doubling time of infB-SPA strain is 37.8 minutes ± 3.3 minutes and MG1655 wild-type is 33.4 minutes ± 1.8 minutes (2-sided t-test of unequal variance P-value is 0.14). Error bars represent the standard deviation in doubling times.

**Fig S8.** Immunofluorescence analysis was performed on wild-type MG1655 to check the specificity of the primary antibody. Since the wild-type MG1655 lacks the SPA-tagged IF-2, no fluorescence signal was detected in these cells, in contrast to the infB-SPA tagged strain, which showed a clear signal. This confirms the specificity of the primary antibody for the SPA-tagged IF-2.

**Fig S9.** Western blot analysis of infB-SPA and BL21 strains using a primary antibody to confirm the conjugation of the SPA tag with the infB gene. The molecular weight of IF-2 is 97 kDa, but due to the presence of the SPA tag, the observed band corresponds to approximately 130 kDa. No band is detected in the BL21 strain, which lacks the SPA tag.

**Fig S10.** List of psychrophilic and thermophilic bacterial species and references for their characterization.

Fig S1

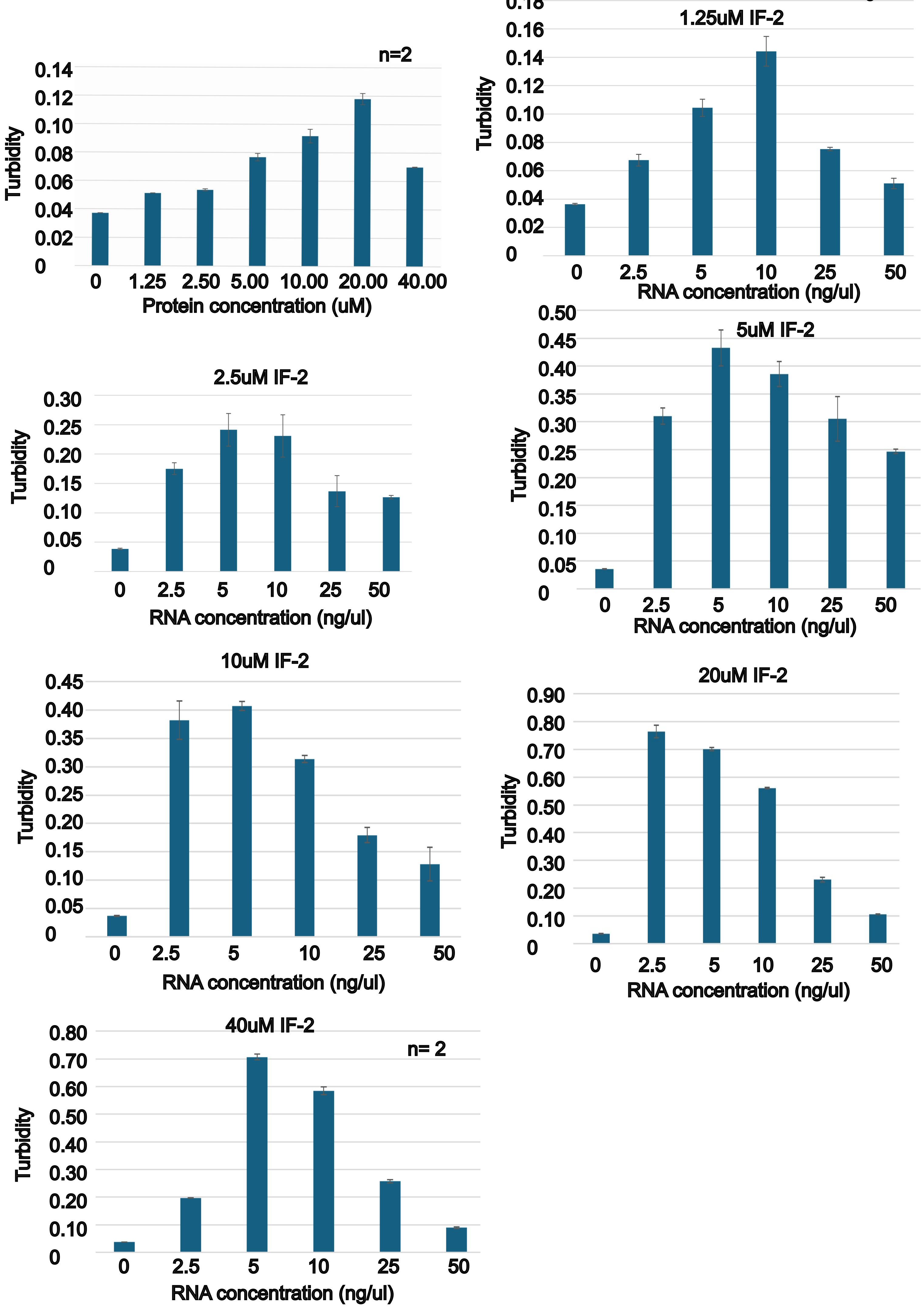
n=2
Turbidity
0.14
0.12
0.10
0.08
0.06
0.04
0.02
0
0
1.25
2.50
5.00
10.00
20.00
40.00
Protein concentration (uM)
1.25uM IF-2
Turbidity
0.18
0.16
0.14
0.12
0.10
0.08
0.06
0.04
0.02
0
0
2.5
5
10
25
50
RNA concentration (ng/ul)
2.5uM IF-2
Turbidity
0.30
0.25
0.20
0.15
0.10
0.05
0
0
2.5
5
10
25
50
RNA concentration (ng/ul)
5uM IF-2
Turbidity
0.50
0.45
0.40
0.35
0.30
0.25
0.20
0.15
0.10
0.05
0
0
2.5
5
10
25
50
RNA concentration (ng/ul)
10uM IF-2
Turbidity
0.45
0.40
0.35
0.30
0.25
0.20
0.15
0.10
0.05
0
0
2.5
5
10
25
50
RNA concentration (ng/ul)
20uM IF-2
Turbidity
0.90
0.80
0.70
0.60
0.50
0.40
0.30
0.20
0.10
0
0
2.5
5
10
25
50
RNA concentration (ng/ul)
40uM IF-2
n= 2
Turbidity
0.80
0.70
0.60
0.50
0.40
0.30
0.20
0.10
0
0
2.5
5
10
25
50
RNA concentration (ng/ul)

Fig S2

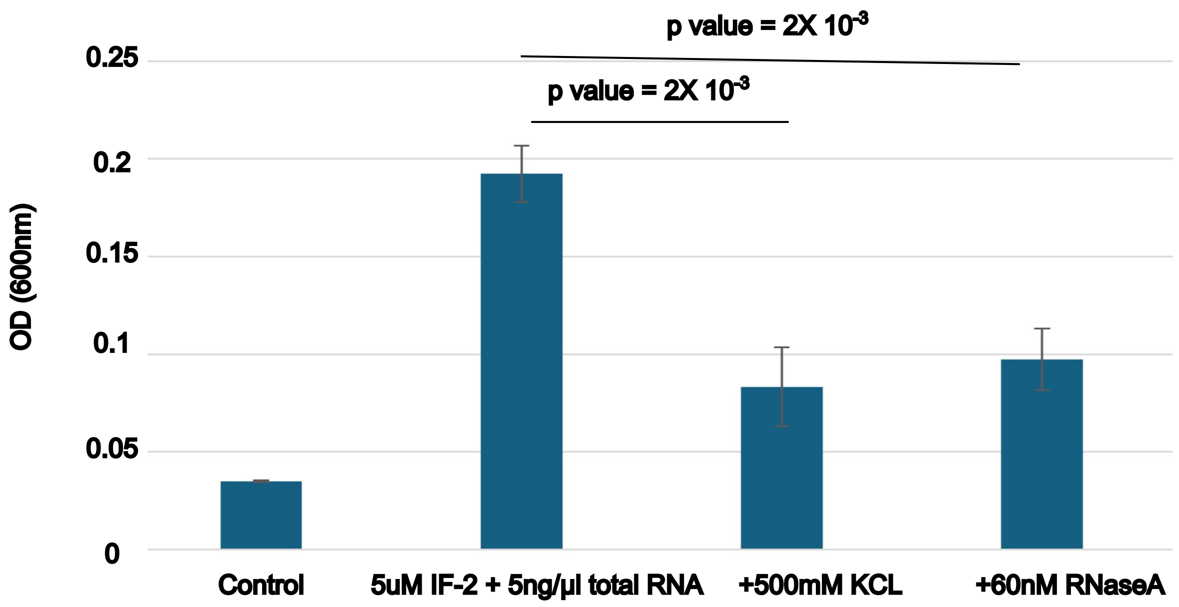
p value = 2X 10-3
p value = 2X 10-3
OD (600nm)
0.25
0.2
0.15
0.1
0.05
0
Control
5uM IF-2 + 5ng/µl total RNA
+500mM KCL
+60nM RNaseA

Fig S3

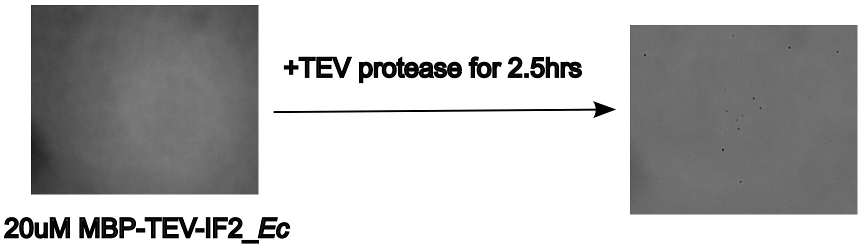

Fig S4

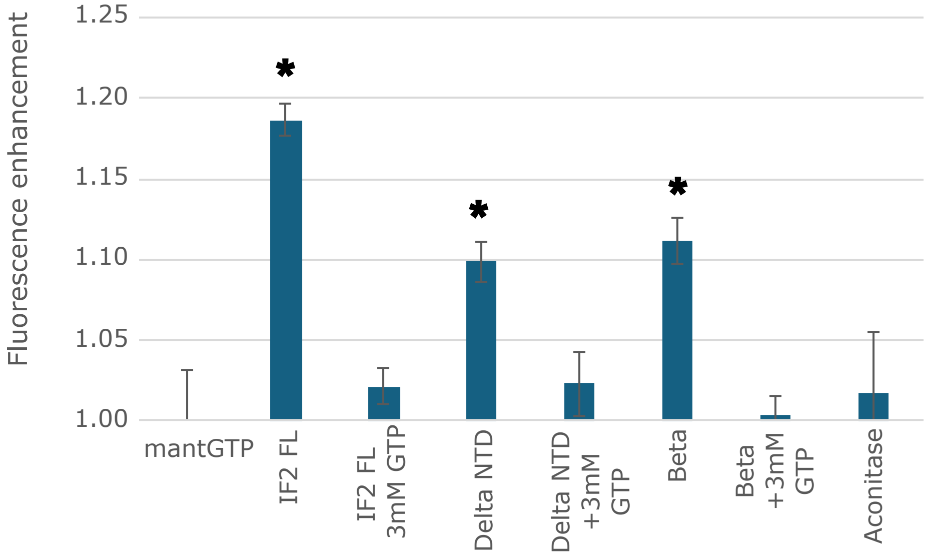
Fluorescence enhancement
1.25
1.20
1.15
1.10
1.05
1.00
*
*
*
mantGTP
IF2 FL
IF2 FL 3mM GTP
Delta NTD
Delta NTD +3mM GTP
Beta
Beta +3mM GTP
Aconitase

Fig S5

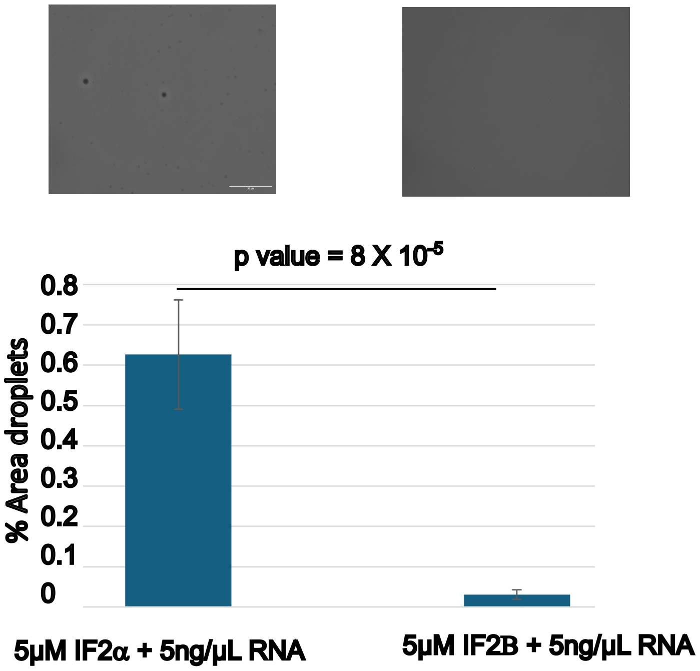
p value = 8 X 10$^{-5}$
% Area droplets
0.8
0.7
0.6
0.5
0.4
0.3
0.2
0.1
0
5μM IF2α + 5ng/μL RNA
5μM IF2B + 5ng/μL RNA

Fig S6

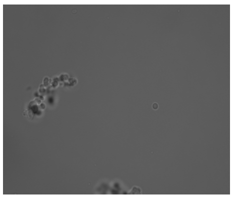

5µM IF-2 + 5ng/µL RNA
+1mM GTP

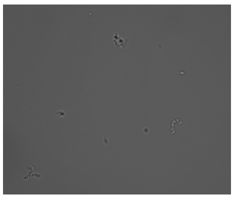

5µM IF-2 + 5ng/µL RNA
+ 1mM GDP

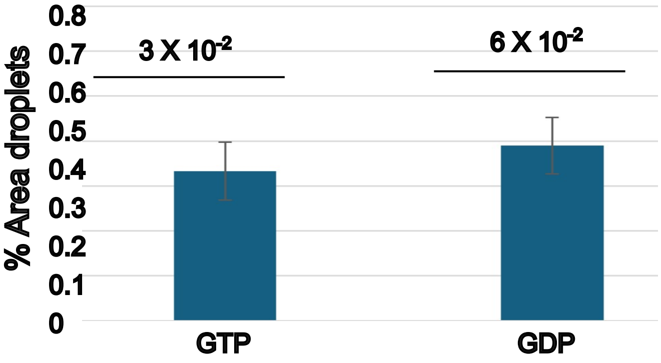

Fig S7

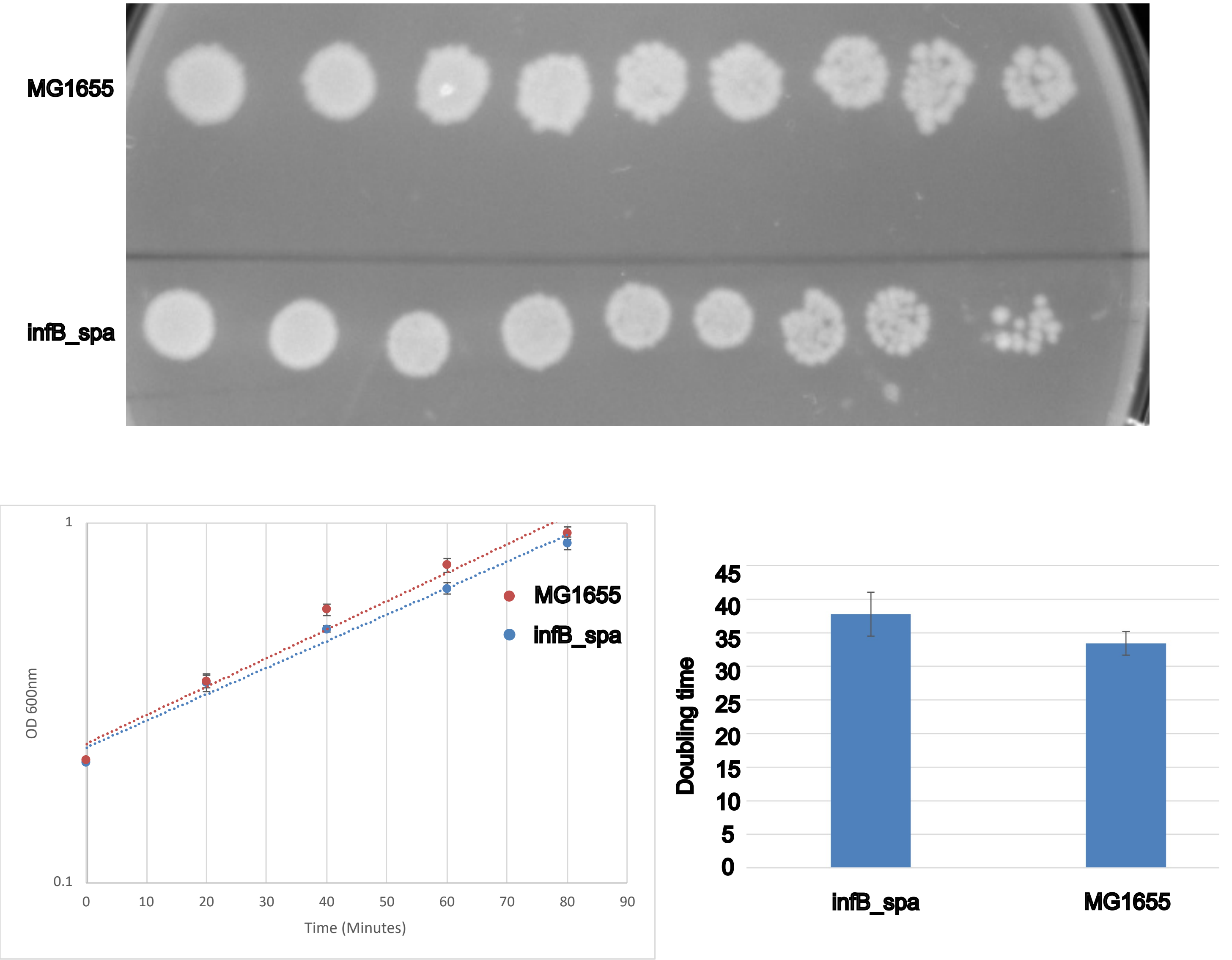
MG1655
infB_spa
1
0.1
0
10
20
30
40
50
60
70
80
90
OD 600nm
Time (Minutes)
MG1655
infB_spa
Doubling time
45
40
35
30
25
20
15
10
5
0
infB_spa
MG1655

Fig S8

**mouse-anti-FLAG primary**
**anti-mouse-CY5 secondary**

MG1655

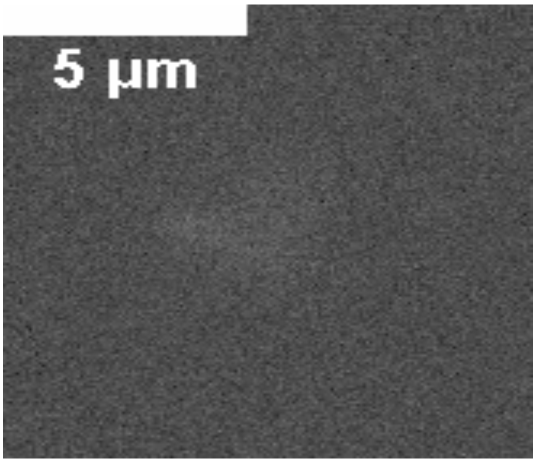


cy5 channel

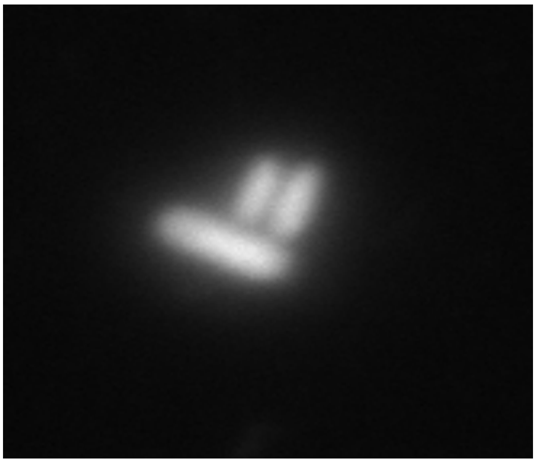

Dapi

infB-SPA

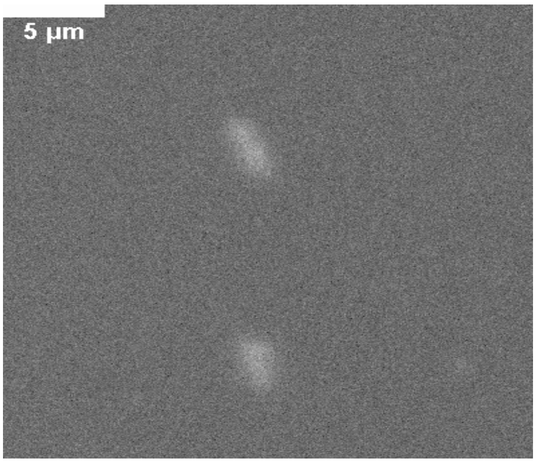


cy5 channel

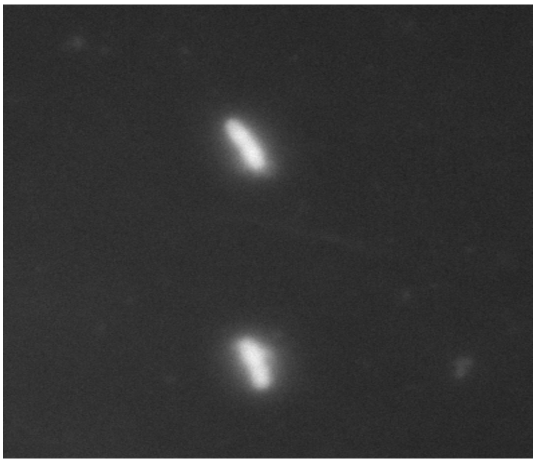

Dapi

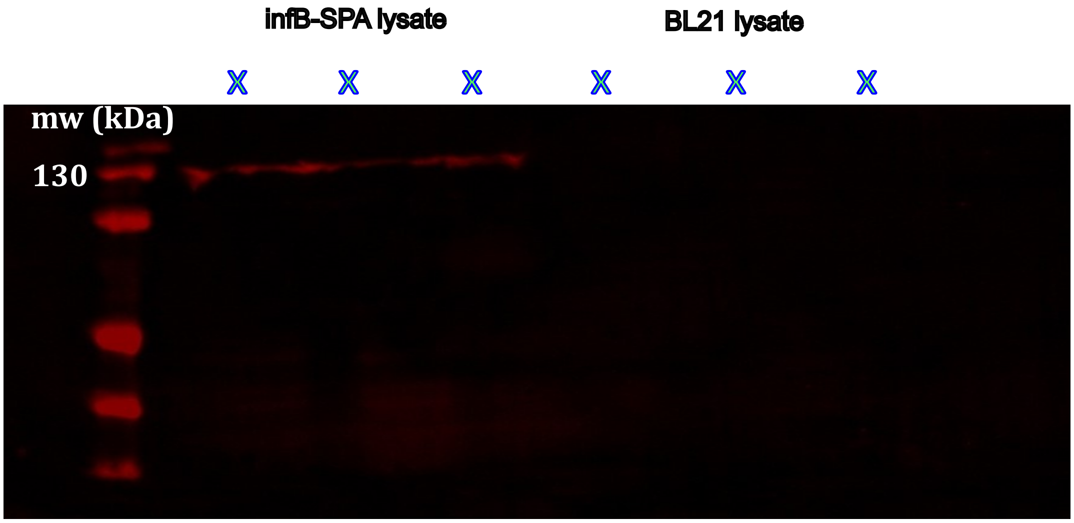
infB-SPA lysate
BL21 lysate
X
X
X
X
X
X
mw (kDa)
130

| Name | Type | Reference |
|---|---|---|
| Pseudorhodobacter antarcticus | Psychrophile | 1 |
| Shewanella livingstonensis | Psychrophile | 2 |
| Psychrobacter cryohalolentis K5 | Psychrophile | 3, 4 |
| Colwellia psychrerythraea 34H | Psychrophile | 3, 5 |
| Psychromonas ingrahamii 37 | Psychrophile | 6 |
| Psychrobacter arcticus 273-4 | Psychrophile | 7, 8 |
| Flavobacterium psychrophilum JIP02/86 | Psychrophile | 9 |
| Desulfotalea psychrophila LSv54 | Psychrophile | 10, 11 |
| Pseudoalteromonas atlantica T6c | Psychrophile | 11 |
| Renibacterium salmoninarum ATCC 33209 | Psychrophile | 11 |
| Octadecabacter arcticus | Psychrophile | 12 |
| Octadecabacter antarcticus | Psychrophile | 12 |
| Thermus thermophilus HB27 | Thermophile | 13 |
| Caldicellulosiruptor saccharolyticus DSM 8903 | Thermophile | 14 |
| Thermus scotoductus SA-01 | Thermophile | 15 |
| Fervidobacterium nodosum Rt17-B1 | Thermophile | 16 |
| Thermodesulfobacterium commune DSM 2178 | Thermophile | 17 |
| Hydrogenobacter thermophilus TK-6 | Thermophile | 18 |
| Thermotoga maritima MSB8 | Thermophile | 19 |
| Elioraea tepidiphila | Thermophile | 20 |
| Tepidamorphus gemmatus | Thermophile | 21 |
| Rubellimicrobium thermophilum | Thermophile | 22 |


(1) Chen, C. X.; Zhang, X. Y.; Liu, C.; Yu, Y.; Liu, A.; Li, G. W.; Li, H.; Chen, X. L.; Chen, B.; Zhou, B. C.; et al. Pseudorhodobacter antarcticus sp. nov., isolated from Antarctic intertidal sandy sediment, and emended description of the genus Pseudorhodobacter Uchino et al. 2002 emend. Jung et al. 2012. *Int J Syst Evol Microbiol* **2013**, *63* (Pt 3), 849-854. DOI: 10.1099/ijs.0.042184-0 From NLM.

(2) Luo, G.; Fujii, S.; Koda, T.; Tajima, T.; Sambongi, Y.; Hida, A.; Kato, J. Unexpectedly high thermostability of an NADP-dependent malic enzyme from a psychrophilic bacterium, Shewanella livingstonensis Ac10. *J Biosci Bioeng* **2021**, *132* (5), 445-450. DOI: 10.1016/j.jbiosc.2021.07.005 From NLM.

(3) Amato, P.; Christner, B. C. Energy metabolism response to low-temperature and frozen conditions in Psychrobacter cryohalolentis. *Appl Environ Microbiol* **2009**, *75* (3), 711-718. DOI: 10.1128/aem.02193-08 From NLM.

(4) Smith, D. J.; Schuerger, A. C.; Davidson, M. M.; Pacala, S. W.; Bakermans, C.; Onstott, T. C. Survivability of Psychrobacter cryohalolentis K5 under simulated martian surface conditions. *Astrobiology* **2009**, *9* (2), 221-228. DOI: 10.1089/ast.2007.0231 From NLM.

(5) Czajka, J. J.; Abernathy, M. H.; Benites, V. T.; Baidoo, E. E. K.; Deming, J. W.; Tang, Y. J. Model metabolic strategy for heterotrophic bacteria in the cold ocean based on Colwellia psychrerythraea 34H. *Proc Natl Acad Sci U S A* **2018**, *115* (49), 12507-12512. DOI: 10.1073/pnas.1807804115 From NLM.

(6) Auman, A. J.; Breezee, J. L.; Gosink, J. J.; Kämpfer, P.; Staley, J. T. Psychromonas ingrahamii sp. nov., a novel gas vacuolate, psychrophilic bacterium isolated from Arctic polar sea ice. *Int J Syst Evol Microbiol* **2006**, *56* (Pt 5), 1001-1007. DOI: 10.1099/ijs.0.64068-0 From NLM.

(7) Ayala-del-Río, H. L.; Chain, P. S.; Grzymski, J. J.; Ponder, M. A.; Ivanova, N.; Bergholz, P. W.; Di Bartolo, G.; Hauser, L.; Land, M.; Bakermans, C.; et al. The genome sequence of Psychrobacter arcticus 273-4, a psychroactive Siberian permafrost bacterium, reveals mechanisms for adaptation to low-temperature growth. *Appl Environ Microbiol* **2010**, *76* (7), 2304-2312. DOI: 10.1128/aem.02101-09 From NLM.

(8) Casillo, A.; Ziaco, M.; Lindner, B.; Parrilli, E.; Schwudke, D.; Holgado, A.; Beyaert, R.; Lanzetta, R.; Tutino, M. L.; Corsaro, M. M. Lipid A structural characterization from the LPS of the Siberian psychro-tolerant Psychrobacter arcticus 273-4 grown at low temperature. *Extremophiles* **2018**, *22* (6), 955-963. DOI: 10.1007/s00792-018-1051-6 From NLM.

(9) Hesami, S.; Metcalf, D. S.; Lumsden, J. S.; Macinnes, J. I. Identification of cold-temperature-regulated genes in Flavobacterium psychrophilum. *Appl Environ Microbiol* **2011**, *77* (5), 1593-1600. DOI: 10.1128/aem.01717-10 From NLM.

(10) Rabus, R.; Ruepp, A.; Frickey, T.; Rattei, T.; Fartmann, B.; Stark, M.; Bauer, M.; Zibat, A.; Lombardot, T.; Becker, I.; et al. The genome of Desulfotalea psychrophila, a sulfate-reducing bacterium from permanently cold Arctic sediments. *Environ Microbiol* **2004**, *6* (9), 887-902. DOI: 10.1111/j.1462-2920.2004.00665.x From NLM.

(11) Metpally, R. P. R.; Reddy, B. V. B. Comparative proteome analysis of psychrophilic versus mesophilic bacterial species: Insights into the molecular basis of cold adaptation of proteins. *BMC Genomics* **2009**, *10* (1), 11. DOI: 10.1186/1471-2164-10-11.

(12) Vollmers, J.; Voget, S.; Dietrich, S.; Gollnow, K.; Smits, M.; Meyer, K.; Brinkhoff, T.; Simon, M.; Daniel, R. Poles apart: Arctic and Antarctic Octadecabacter strains share high genome plasticity and a new type of xanthorhodopsin. *PLoS One* **2013**, *8* (5), e63422. DOI: 10.1371/journal.pone.0063422 From NLM.

(13) Kirchner, L.; Müller, V.; Averhoff, B. A temperature dependent pilin promoter for production of thermostable enzymes in Thermus thermophilus. *Microb Cell Fact* **2023**, *22* (1), 187. DOI: 10.1186/s12934-023-02192-1 From NLM.

(14) Talluri, S.; Raj, S. M.; Christopher, L. P. Consolidated bioprocessing of untreated switchgrass to hydrogen by the extreme thermophile Caldicellulosiruptor saccharolyticus DSM 8903. *Bioresour Technol* **2013**, *139*, 272-279. DOI: 10.1016/j.biortech.2013.04.005 From NLM.

(15) Cockrell, A. L.; Fitzgerald, L. A.; Cusick, K. D.; Barlow, D. E.; Tsoi, S. D.; Soto, C. M.; Baldwin, J. W.; Dale, J. R.; Morris, R. E.; Little, B. J.; et al. Differences in Physical and Biochemical Properties of Thermus scotoductus SA-01 Cultured with Dielectric or Convection Heating. *Appl Environ Microbiol* **2015**, *81* (18), 6285-6293. DOI: 10.1128/aem.01618-15 From NLM.

(16) Yang, Y.; Zhu, Y.; Obaroakpo, J. U.; Zhang, S.; Lu, J.; Yang, L.; Ni, D.; Pang, X.; Lv, J. Identification of a novel type I pullulanase from Fervidobacterium nodosum Rt17-B1, with high thermostability and suitable optimal pH. *Int J Biol Macromol* **2020**, *143*, 424-433. DOI: 10.1016/j.ijbiomac.2019.10.112 From NLM.

(17) Bhatnagar, S.; Badger, J. H.; Madupu, R.; Khouri, H. M.; O'Connor, E. M.; Robb, F. T.; Ward, N. L.; Eisen, J. A. Genome Sequence of a Sulfate-Reducing Thermophilic Bacterium, Thermodesulfobacterium commune DSM 2178T (Phylum Thermodesulfobacteria). *Genome Announc* **2015**, *3* (1). DOI: 10.1128/genomeA.01490-14 From NLM.

(18) Ishii, M.; Igarashi, Y.; Kodama, T. Purification and characterization of ATP:citrate lyase from Hydrogenobacter thermophilus TK-6. *J Bacteriol* **1989**, *171* (4), 1788-1792. DOI: 10.1128/jb.171.4.1788-1792.1989 From NLM.

(19) Wang, Z.; Tong, W.; Wang, Q.; Bai, X.; Chen, Z.; Zhao, J.; Xu, N.; Liu, S. The temperature dependent proteomic analysis of Thermotoga maritima. *PLoS One* **2012**, *7* (10), e46463. DOI: 10.1371/journal.pone.0046463 From NLM.

(20) Saini, M. K.; Yoshida, S.; Sebastian, A.; Hara, E.; Tamaki, H.; Soulier, N. T.; Albert, I.; Hanada, S.; Tank, M.; Bryant, D. A. Elioraea tepida, sp. nov., a Moderately Thermophilic Aerobic Anoxygenic Phototrophic Bacterium Isolated from the Mat Community of an Alkaline Siliceous Hot Spring in Yellowstone National Park, WY, USA. *Microorganisms* **2022**, *10* (1), 80.

(21) Albuquerque, L.; Rainey, F. A.; Pena, A.; Tiago, I.; Veríssimo, A.; Nobre, M. F.; da Costa, M. S. Tepidamorphus gemmatus gen. nov., sp. nov., a slightly thermophilic member of the Alphaproteobacteria. *Syst Appl Microbiol* **2010**, *33* (2), 60-66. DOI: 10.1016/j.syapm.2010.01.002 From NLM.

(22) Denner, E. B. M.; Kolari, M.; Hoornstra, D.; Tsitko, I.; Kämpfer, P.; Busse, H. J.; Salkinoja-Salonen, M. Rubellimicrobium thermophilum gen. nov., sp. nov., a red-pigmented, moderately thermophilic bacterium isolated from coloured slime deposits in paper machines. *Int J Syst Evol Microbiol* **2006**, *56* (Pt 6), 1355-1362. DOI: 10.1099/ijs.0.63751-0 From NLM.

(23) Altschul, S. F.; Gish, W.; Miller, W.; Myers, E. W.; Lipman, D. J. Basic local alignment search tool. *J Mol Biol* **1990**, *215* (3), 403-410. DOI: 10.1016/s0022-2836(05)80360-2 From NLM.

(24) Sayers, E. W.; Bolton, E. E.; Brister, J. R.; Canese, K.; Chan, J.; Comeau, D. C.; Connor, R.; Funk, K.; Kelly, C.; Kim, S.; et al. Database resources of the national center for biotechnology information. *Nucleic Acids Res* **2022**, *50* (D1), D20-d26. DOI: 10.1093/nar/gkab1112 From NLM.

(25) Emenecker, R. J.; Griffith, D.; Holehouse, A. S. Metapredict: a fast, accurate, and easy-to-use predictor of consensus disorder and structure. *Biophysical Journal* **2021**, *120* (20), 4312-4319. DOI: https://doi.org/10.1016/j.bpj.2021.08.039.

(26) Gasteiger E., H. C., Gattiker A., Duvaud S., Wilkins M.R., Appel R.D., Bairoch A. Protein Identification and Analysis Tools on the Expasy Server. Walker, J. M. Ed.; The Proteomics Protocols Handbook, Humana Press, 2005; pp 571-607.
